# Neurophysiological signatures of default mode network dysfunction and cognitive decline in Alzheimer’s disease

**DOI:** 10.1101/2024.09.16.613373

**Authors:** Recep A. Ozdemir, Brice Passera, Peter J. Fried, Daniel Press, Lynn W. Shaughnessy, Stephanie Buss, Mouhsin M. Shafi

## Abstract

Neural hyper-excitability and network dysfunction are neurophysiological hallmarks of Alzheimer’s disease (AD) in animal studies, but their presence and clinical relevance in humans remain poorly understood. We introduce a novel perturbation-based approach combining transcranial magnetic stimulation and electroencephalography (TMS-EEG), alongside resting-state EEG (rsEEG), to investigate neurophysiological basis of default mode network (DMN) dysfunction in early AD. While rsEEG revealed global neural slowing and disrupted synchrony, these measures reflected widespread changes in brain neurophysiology without network-specific insights. In contrast, TMS-EEG identified network-specific local hyper-excitability in the parietal DMN and disrupted connectivity with frontal DMN regions, which uniquely predicted distinct cognitive impairments and mediated the link between structural brain integrity and cognition. Our findings provide mechanistic insights into how network-specific neurophysiological disruptions contribute to AD-related cognitive dysfunction. Perturbation-based assessments hold promise as novel markers of early detection, disease progression, and target engagement for disease-modifying therapies aiming to restore abnormal neurophysiology in AD.

## Introduction

Alzheimer’s disease (AD) is marked by the early deposition of amyloid-beta-42 (Aβ) plaques followed by the rapid accumulation of phosphorylated tau (p-tau) in neurofibrillary tangles^1–3^, and is associated with significant neurophysiological changes before the onset of clinical symptoms^4^. In animal models, Aβ plaques disrupt synaptic activity, causing an imbalance in inhibition and excitation, with neurons surrounding these plaques becoming abnormally hyperactive^5^. P-tau independently suppresses neural activity^6^, leading to synaptic loss, and significant cerebral atrophy^7^ in brain regions critical for memory function^2^. Advanced imaging methods like positron emission tomography (PET) have transformed the monitoring of these pathological markers in AD patients, enhancing our understanding of their relevance to disease progression. However, the neurophysiological manifestations of AD pathology and their clinical significance, remain poorly understood in humans.

Neural activity and connectivity dynamics in AD have been largely investigated using resting-state functional MRI (rsfMRI), with the default mode network (DMN) showing early and significant disruptions^8,9^. Early rsfMRI studies reported decreased activity in the posterior cingulate cortex (PCC) and inferior parietal lobule (IPL)^10–12^, findings that align with PET studies showing increased amyloid deposition^13^ and reduced metabolic activity in these regions^14,15^. Recent research has identified the posterior DMN as the first region to exhibit abnormal connectivity patterns that correlate with cognitive decline^16^ and neuropathology^17^ across the AD spectrum. While this research has greatly advanced our understanding of the functional network organization in AD pathology, rsfMRI primarily captures correlational changes in blood oxygenation across the brain, providing indirect estimates of neuronal activity and network connectivity with limited capacity to detect synaptic disruptions at the individual level^18,19^. High temporal resolution methods such as electroencephalography (EEG) and magnetoencephalography (MEG) are therefore preferred for measuring synchronized synaptic activity across large-scale brain networks. In AD patients, these neurophysiological modalities have consistently shown a global slowing of spectral power dynamics, with increased power in slower and decreased power in faster neural oscillations^20–26^. MEG connectivity analyses generally show decreased alpha (8-12 Hz) over visual cortices and increased delta-theta (1-7 Hz) neural synchrony across the cortex^27,28^. One limitation of these resting state based EEG/MEG measures is that they do not directly reflect AD-related neural hyper-excitability and network dysconnectivity. Spontaneous neural oscillations may not be optimal for localizing network-specific neurophysiological dynamics, as neural activity during the resting state arises from multiple sources^29^ with a diffuse and widespread distribution across overlapping networks^30^. Additionally, without external stimuli, spectral power estimates can only serve as surrogates for cortical excitability.

Combining transcranial magnetic stimulation (TMS) with EEG offers a promising approach to address some limitations of both rsfMRI and rsEEG. Single pulses of TMS (spTMS) initially excite the axonal terminals of cortical pyramidal and interneurons^31^, particularly at gyral crowns of the cortex, and evoke a series of high-frequency synaptic activations that directly represent local cortical excitability at the site of stimulation^32–35^. These early activations are followed by trans-synaptic network responses, reflecting the connectivity profile of the stimulated brain region^36^. EEG effectively captures synchronous post-synaptic potentials in pyramidal and interneurons with precise temporal resolution and has the highest sensitivity to neural activity at the superficial layers of the cortex^37^, making it an ideal neurophysiological modality to index the activity of neural populations targeted by TMS. In AD, TMS can directly stimulate well-defined nodes of the DMN to assess local activation and connectivity abnormalities. However, most TMS research in AD has focused on the motor cortex^38–46^, with only a few recent studies examining other brain regions^47–50^. These recent TMS-EEG studies have typically performed electrode-level analyses at local TMS sites, offering a limited understanding of the functional network dynamics implicated in AD. Recently, we developed an MRI-guided TMS-EEG method that personalizes the topography of fMRI-based functional networks on individual brains, identifies network-specific TMS targets, and generates local activation and causal network connectivity maps at the individual level^51^. Using this approach, we demonstrated that EEG responses to TMS provide an accurate ‘fingerprint’ of individual brain activation patterns^52^, preferentially propagate through the structural connectivity of the stimulated network^53,54^, characterize causal network connectivity dynamics with high reproducibility^55,56^, and identify brain-behavior relationships not observable through resting-state recordings in healthy individuals^51^.

In this study, we aimed to characterize the neurophysiological signatures of the DMN dysfunction in AD by utilizing both TMS-EEG and rsEEG measures, and evaluated their relationships to cognition. Our study is the first to comprehensively integrate rsEEG and TMS-EEG within the same cohort, offering a unique opportunity to compare these modalities and uncover network-specific abnormalities in AD. Using spTMS, we targeted individually defined regions within the left IPL (IPL-TMS), a DMN node particularly vulnerable to AD pathology, in biomarker-confirmed AD patients (*n*=37) and age-matched cognitively normal controls (*n*=40). Additionally, we stimulated the left primary motor cortex (M1-TMS) as a control site and included a sham TMS condition (Sham-TMS). RsEEG was recorded for five minutes during an eyes-closed condition to assess spontaneous neural dynamics. We further examined how these measures relate to specific cognitive impairments, accounting for participant demographics (age, education) and structural brain integrity at TMS targeted brain regions. While we expect spectral slowing and reduced alpha neural synchrony in AD patients, we hypothesize that rsEEG measures will reflect global brain changes but will not be specific to the DMN. In contrast, TMS-EEG measures would reveal DMN-specific dysfunction in AD, characterized by hyper-excitability in the IPL and abnormal hyper- and hypo-connectivity patterns with non-stimulated DMN regions, providing insight into the network-specific mechanisms underlying decline in distinct cognitive functions.

## Results

### TMS of DMN evokes distinct spatial-temporal neural activation dynamics in AD

We performed source space reconstruction using individual MRIs and digitized electrode locations to measure TMS-evoked (perturbation-based) cortical excitability and network connectivity (Fig. 1). We generated local cortical maps based on electric field (E-field) simulations of personalized TMS targets^57,58^, where spTMS would most likely to generate direct neural responses^31^, to measure cortical excitability at each TMS target with high spatial accuracy (Fig. 1B). For network connectivity, we morphed canonical DMN maps^59^ onto individual brains (Fig. 1C) and measure how much of the TMS evoked activity propagated from local (inferior parietal) to non-stimulated DMN nodes (Fig. 1D).

**Fig. 1.**
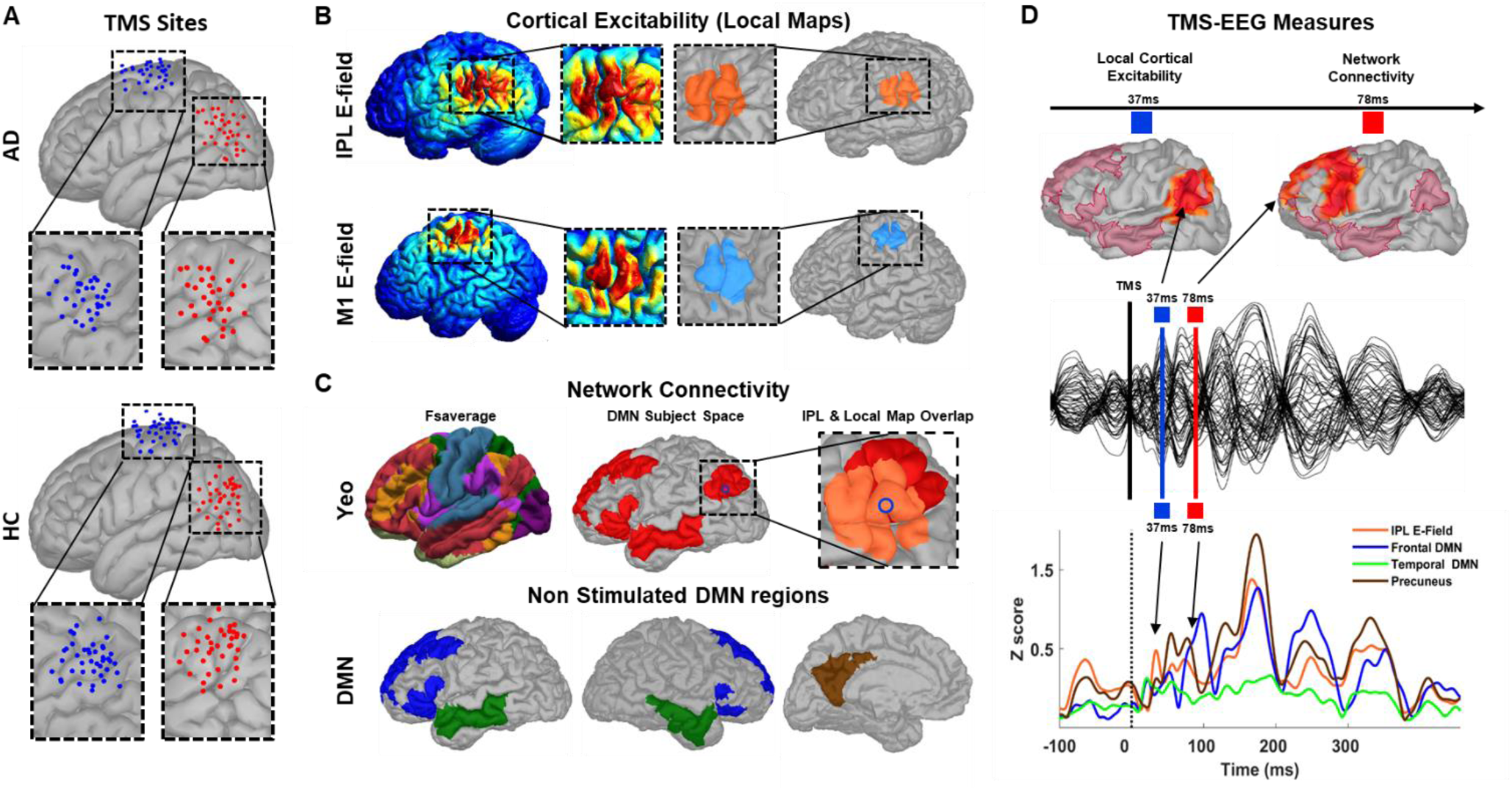
TMS Targets and Overview of TMS-EEG Measure Extraction in Source Space. **A:** Individual TMS targets for AD (upper) and HC (lower) groups projected on a common brain template for each stimulation site (M1: blue dots, IPL: red dots). **B:** Individual E-field simulation maps (left panels) were thresholded at the highest % 0.01 to generate personalized local masks (right panels) on cortical surface for each TMS site (upper panel for IPL and lower panel for M1). These local masks were used to extract average regional EEG activity and measure cortical excitability at the site of stimulation. **C:** Canonical rsfMRI maps derived from group-averaged fMRI connectivity analysis (left) were projected onto individual cortical surfaces and DMN mask (middle panel) used as region of interests. TMS targeted DMN node (IPL) is highlighted within the squared region with the blue dot representing TMS site (middle). This region is expanded on the right to show overlap between IPL-DMN (red) and E-field based local map (orange). Lower panel shows distinct non-stimulated regions at frontal (blue), temporal (green) and precuneus (brown) sites within the DMN mask. These masks are used to measure network connectivity. **D:** Representative example of TMS-EEG measures. Upper panel shows individual cortical surface model with DMN mask (shaded red regions) and source reconstructed EEG activity for IPL-TMS from a representative HC participant. Thresholded EEG activity (> %70) at 37ms following TMS is localized over the stimulated region (Left-IPL) representing local cortical excitability (left), while EEG activity at 79ms (right) is localized over frontal DMN representing network connectivity. Middle panel shows scalp TEPs used for source reconstruction. Lower panel shows time series of normalized averaged EEG activity (in z-scores) extracted from local masks for cortical excitability (orange) and non-stimulated DMN masks for network connectivity measures.

In a representative HC participant, IPL-TMS evoked an early response localized to the stimulation site, with activity propagating to frontal brain regions around 110ms, aligning with the DMN map (Fig. 2A). In an AD participant, IPL-TMS induced high-amplitude early responses localized to the stimulation site, with subsequent activations confined to temporal and posterior regions with restricted propagation to frontal-DMN regions (Fig. 2A). These activation dynamics were specific to the stimulated region and significantly larger than those from Sham-TMS (Fig. 2B).

**Fig. 2.**
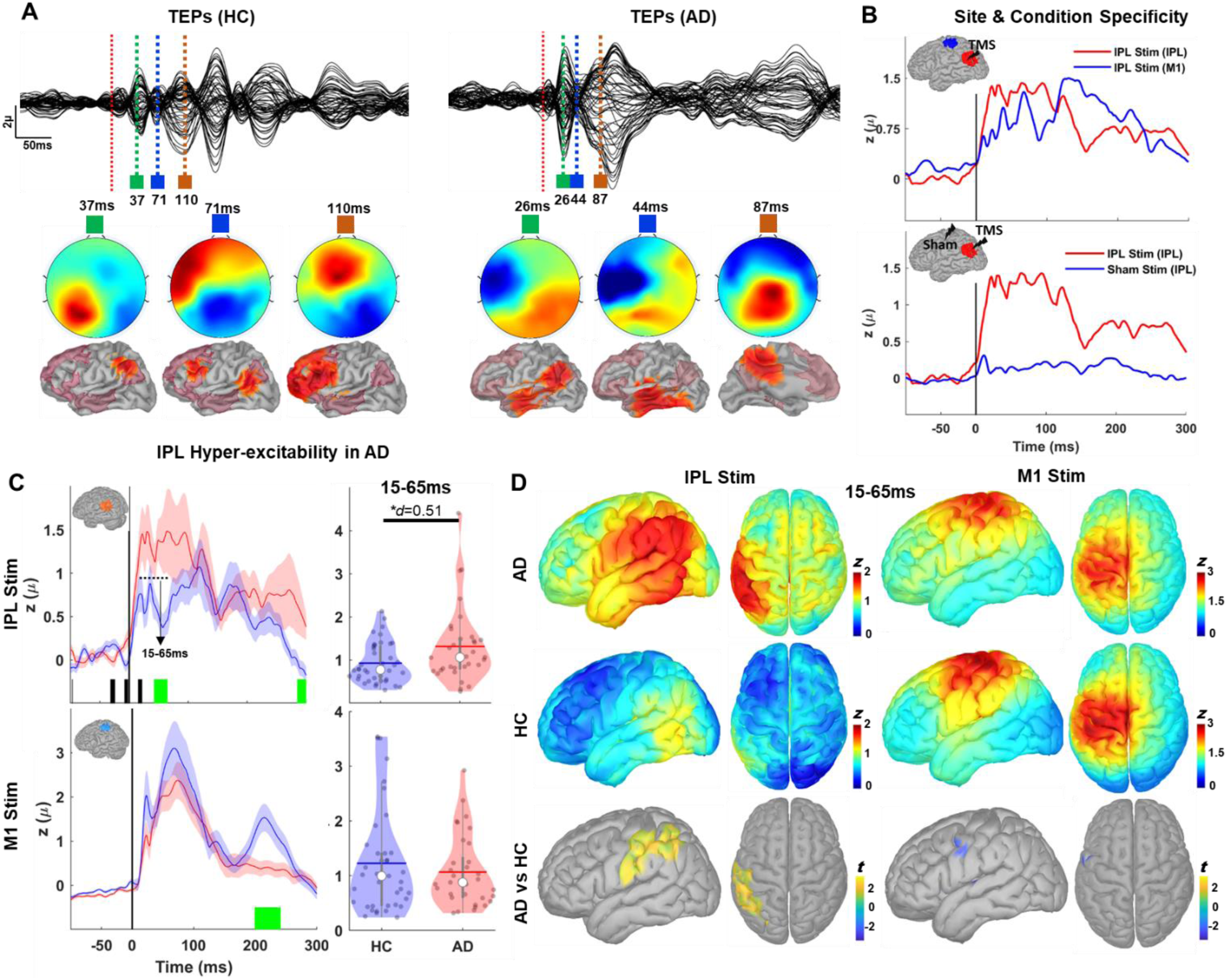
Increased Early-Local DMN Excitability in AD. **A:** Illustration of TMS evoked potentials (TEPs) and their spatial-temporal dynamics from one representative HC (left panels) and one AD participant (right panels). Upper panels show TEPs of all EEG channels after preprocessing with red dotted vertical lines indicating TMS time points and following colored vertical lines showing selected TEP peaks. Middle and lower panels show corresponding scalp topographies and source reconstructed activations for the selected peaks. **B:** Average current density time series in the AD group showing site and condition specificity of TEPs. Upper panel compares evoked responses derived from IPL (red) and M1 (blue) local masks following stimulation of IPL while lower panel compares IPL responses following IPL (red) and Sham (blue) stimulation. Sham responses were presented after removing auditory evoked potentials. Black vertical lines in each panel indicates TMS time points and brain figures with colored masks show signal extraction and stimulation sites in each condition. **C:** Averaged evoked responses following IPL-TMS (upper left) and M1-TMS (lower left) at the site of stimulation both for AD (red) and HC (Blue) groups. Solid colored lines show group averaged current density time series (in z-scores) extracted from individualized e-field based local masks for IPL (orange colored shared areas of the representative cortical surface in the upper panel) and M1 (light blue shaded areas of the cortical surface in the lower panel). Shaded regions show inter-individual response variation with standard error of measurements (SE). Green colored blocks at the bottom of each panel show significant time-points between AD and HC groups that survived cluster permutations while black colored blocks indicate significant time points that did not survive permutation tests. Violin plots on the right panels show average evoked current densities between 15-65ms following IPL-TMS (designated with black dotted line and arrow on upper right panel) and M1 (lower right) for both groups. White dots in violin plots represent median value and gray colored dots represent individual responses. Colored horizontal lines and gray vertical bars represent grand average values and interquartile ranges, respectively. * in upper violin plot denotes statistical significance at *p*<0.05 with corresponding effect size calculated using Cohens’ *d*. **D:** Cortical maps for grand averaged current densities (in z-scores) between 15-65ms on MNI template following IPL-TMS (left) and M1-TMS (right) for AD (upper panels) and HC (middle panels) groups. Lower panels show statistical results of thresholded cluster-based permutation t-tests (cluster *p*<0.05) with hot colors indicating AD > HC and cold colors indicating AD < HC.

### Local DMN hyper-excitability in AD

We compared local IPL responses in the temporal domain using current density time series from individual brain models (Fig. 2C). IPL-TMS evoked larger local responses between 15 and 65ms in the AD group (Fig. 2C, left upper panel) with two distinct clusters: first from 15-to-22ms (t_(70)_=2.12, *p*=0.039) and second from 42-to-65ms (t_(70)_=2.76, *p*=0.012) with permutations analyses revealed significance for the second cluster (Permutation *p* = 0.034). We also observed a late cluster (t_(70)_=2.17, *p*=0.029) with larger AD responses starting at 285ms following TMS (Permutation *p*=0.042), indicating extended IPL activation in AD. While no significant early response differences were found between the two groups following M1-TMS, HC participants had significantly larger responses that survived permutations (Permutation *p*=0.009) between 199-240ms (Fig. 2C). As early response differences were confined to 15-65ms, we extracted average responses within this time window for each individual for further cortical excitability analyses. Group average early activations were significantly higher in the AD group compared to HC (t_(70)_=2.24, *p*=0.028, *d*=0.51) for IPL stimulation. To further explore the spatial characteristics of early response differences, we averaged whole-brain source activations at the individual level and projected them to a common brain template (Montreal Neurological Institute “MNI”) to generate group-level cortical activation maps (Fig. 2D). IPL-TMS revealed higher activation in the AD primarily centered in the left parietal cortex, involving the inferior and superior parietal lobules, and extending into the postcentral and angular gyri (Fig. 2D). Stimulation of M1 revealed lower activation in the AD group over a small portion of left inferior precentral gyrus but this cluster did not survive permutation. To test the specificity of these results, we compared early IPL responses (15-65ms) following M1 and Sham-TMS (Supplementary Fig. 1A and B). No significant IPL response differences were found following M1 or Sham-TMS, suggesting that cortical hyper-excitability in the AD group is specific to IPL stimulation.

### Hyper-connectivity in Temporal DMN and Hypo-connectivity in Frontal DMN in AD

We extracted average EEG activity from non-stimulated DMN regions following IPL-TMS to assess differences in response propagation between the groups (Fig. 3). AD participants showed greater responses in the left temporal-DMN between 60-85ms, with a significant cluster (t_(70)_=2.37, *p*=0.022) within the 64-78ms period (Permutation *p*=0.026). However, AD participants exhibited smaller activations at the frontal-DMN at two distinct time windows: first window ranging from 75-to-101ms and second ranging from 125-to-175ms. Permutation analyses for these windows showed significant clusters in the 81-96ms (t_(70)_=-2.93, *p*=0.014, Permutation *p*=0.021) and 147-165ms (t_(70)_=-2.41, *p*=0.017, Permutation *p*=0.022) periods. Group average activity in the temporal-DMN (*t*_(70)_=2.51, *p*=0.014) was higher in the AD group for 60-85ms window, but lower in the frontal-DMN both for 75-101ms (*t*_(70)_=-2.91, *p*=0.022) and 125-175ms windows (*t*_(70)_=-3.23, *p*=0.010, Supplementary Fig.2A), suggesting a selective increase in temporal-DMN connectivity but decreased connectivity to frontal-DMN following IPL-TMS. No significant differences were found in the precuneus following IPL-TMS (*p* > 0.05, data not shown). Group-level cortical activation maps (Fig. 3B) revealed higher activity in AD primarily over the left temporal and inferior parietal cortices but lower activity over the premotor dorsolateral prefrontal and superior frontal cortices. No significant group differences were observed for sham-TMS (Supplementary Fig. 3).

**Fig. 3.**
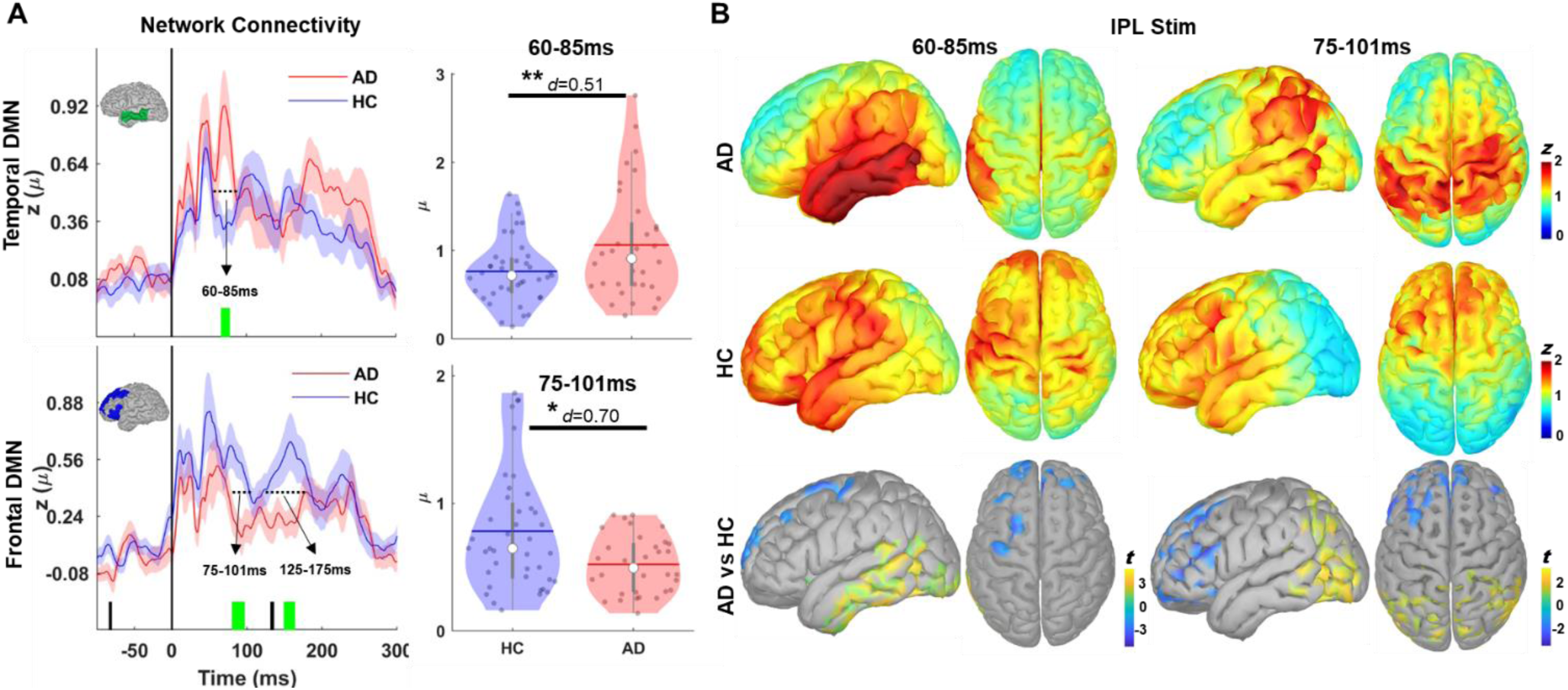
Increased Temporal and Reduced Frontal DMN Connectivity in AD. **A:** Temporal dynamics of TMS evoked responses at temporal (green colored brain regions on upper left panel) and frontal DMN (dark blue colored brain region on lower left panel) following IPL stimulation. Solid colored lines show group averaged current density time series (in z-scores) while shaded regions showed variation with SE of measurements. Green Colored blocks at the bottom of each panel show significant cluster of time-points between AD and HC groups. Violin plots on the right panels show total amount of evoked current densities between 60-85ms (upper panel) and 75-101ms (lower panel) following IPL-TMS for temporal and frontal DMN, respectively. Violin plots for 125-175 time window is provided in Supplementary Fig. 2. **B:** Cortical maps of grand averaged current densities (in z-scores) between 60-85ms (left) and 75-101ms (right) on a template brain model following IPL-TMS for AD (upper panels) and HC (middle panels) groups. Lower panels show statistical results of thresholded cluster-based permutation t-tests (cluster *p*<0.05) with hot colors indicating AD > HC and cold colors in indicating AD < HC. Cortical maps for 125-175 time window is provided in Supplementary Fig. 2.

### Local hyper-excitability, increased parietal-temporal and reduced parietal-frontal connectivity correlate to poor cognition in AD

We explored how local cortical excitability and network connectivity abnormalities relate to global and specific cognitive functions in the AD group (Fig. 4). We used the Alzheimer’s disease Assessment Scale-Cognitive Subscale (ADAS-cog) for global cognition, the Rey Auditory Verbal Learning Test (RAVLT-Total) for verbal memory, the Digit-Span Backward task for working memory, and the Animal Fluency test for semantic memory and executive function assessments (Supplementary Table. 1). Bivariate correlations were run between potential confounders (age, education, cortical thickness) and our variables of interest (cortical excitability, network connectivity, ADAS-Cog, RAVLT-Total, Digit-Span, and Fluency) (Supplementary Table. 2). Cortical thickness at IPL was moderately correlated with better working memory (Digit-Span: *r*=0.53, *p*=0.004), semantic memory (Fluency: *r*=0.46, *p*=0.015), and global cognition (ADAS-Cog: *r*=-0.39, *p*=0.037). IPL cortical thickness also correlated moderately with increased frontal-DMN connectivity at 75-101ms (*r*=0.47, *p*=0.010). Age was moderately correlated with increased cortical excitability (*r*=0.36, *p*=0.03) and poor memory (RAVLT-Total: *r*=-0.37, *p*=0.03), while education showed no meaningful correlations.

**Fig. 4.**
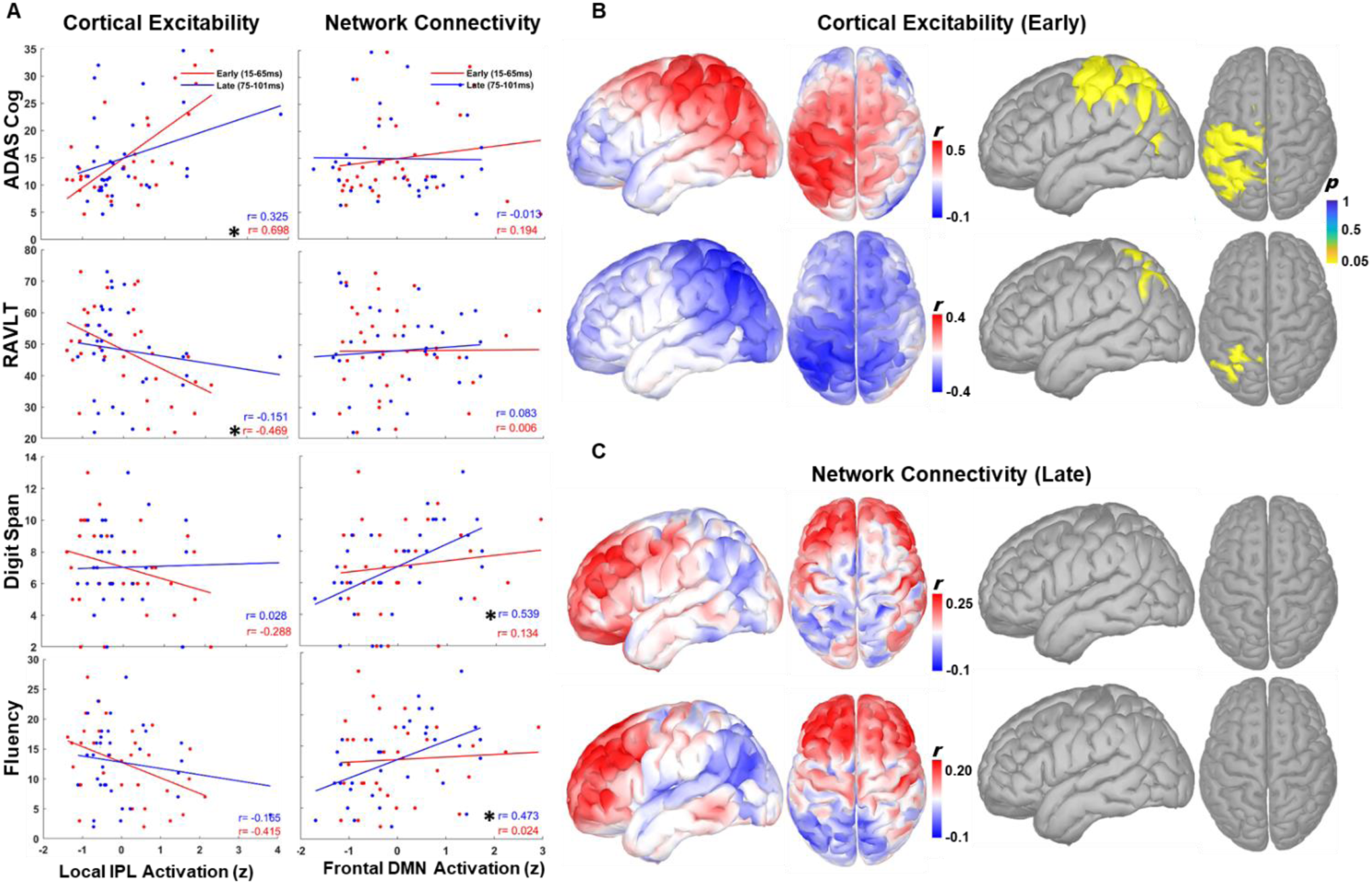
Relationships between TMS-EEG measures of cortical excitability, network connectivity and cognitive functions in AD participants. **A:** Scatter plots with regression lines showing bivariate correlations of cortical excitability (left panels) and network connectivity (right panels) with cognitive function. Color codes refer different temporal windows with red showing early (15-65ms) and blue showing late (75-101ms) activations. Correlation coefficients are provided for each regression line with asterisks indicating statistically significant correlations (*p*<0.05). **B:** Cortical maps showing vertex-wise correlations of early responses with ADAS-cog (upper left) and RAVLT (lower left) scores with hot colors indicating positive and cold colors indicating negative correlations. Statistically thresholded cortical maps (*p*<0.05) on the right panel shows brain regions with significant correlations. **C:** Cortical maps showing vertex-wise correlations of late responses with Digit-Span (Upper panels) and Fluency (Lower panels).

IPL cortical hyper-excitability was strongly correlated with poor global cognition (ADAS-Cog: *r*=0.70, *p*<0.001), moderately with poor verbal memory (RAVLT-Total: *r*=-0.47, *p*=0.01) and executive functions (Fluency: *r*=-0.41, *p*=0.01) (Fig. 4A). Hierarchical regression analyses controlling for age, education, and IPL cortical-thickness showed that IPL hyper-excitability significantly predicted global cognition, explaining an additional 34.3% of unique variance in ADAS-Cog scores (*β*=0.641, *t*=4.061, Δ*R*²=0.34, *p*<0.001). When controlling for cortical excitability, however, cortical thickness was no longer a significant predictor of ADAS-Cog (*p*>0.05), suggesting that hyper-excitability may partially mediate this relationship (Supplementary Fig. 4A). Notably, we noted a temporal specificity between IPL hyper-excitability and cognition such that only early responses between 15-65ms period were strongly correlated with global cognition and moderately correlated with memory functions (red scatter plots and regression lines in Fig. 4A left panels), while late responses between 75-101ms showed zero correlations. Correlations with local M1 responses following M1-TMS and local IPL responses following Sham-TMS were not significant, suggesting site-specificity of these correlations (Supplementary Fig. 5A). Similar patterns were seen for temporal and precuneus DMN, except for a selective increase in the negative relationship between late temporal-DMN activity (75-101ms) and RAVLT suggesting that hyper-connectivity in temporal-DMN is associated with poor verbal memory (Supplementary Fig. 6).

For frontal DMN connectivity, late responses at 75-101ms window were moderately correlated with better working (Digit-Span Backward: *r*=0.54, *p*=0.01) and semantic memory executive (Fluency: *r*=0.47, *p*=0.01) functions (Fig. 5A right panels). Hierarchical regression analyses controlling for age, education and IPL cortical-thickness, showed that frontal-DMN connectivity significantly predicted better Digit-Span (*β*=0.405, *t*=2.234, Δ*R*²=0.107, *p*=0.036) and Fluency (*β*=0.512, *t*=3.043, Δ*R*²=0.226, *p*=0.006) performance, explaining an additional 10% and 22% of unique variance, respectively. After controlling for frontal-DMN connectivity, IPL cortical-thickness remained a significant predictor for Digit-Span (*β*=0.382, *t*=2.084, Δ*R*²=0.903, *p*=0.048) but not for Fluency (*p*>0.05, suggesting a mediation effect of network connectivity for the relationship between cortical-thickness and fluency (Supplementary Fig. 4B). Similar relationship pattern was also observed for late responses at 125-175ms window with frontal-DMN connectivity being significant predictor of better working and semantic memory performance (Supplementary Fig.2C). Frontal-DMN and cognition correlations were specific to IPL stimulation, as M1- and Sham-TMS frontal responses showed no significant correlations (Supplementary Fig. 5B). Finally, vertex-based correlation analyses using showed that increased early IPL excitability (15-65ms) was significantly related to poorer global cognition and memory in the left parietal cortex for ADAS-Cog and angular gyrus for RAVLT-Total (Fig. 4B). Increased signal propagation to frontal regions following IPL stimulation was associated with better working and semantic memory functions (Fig. 4C).

**Fig. 5.**
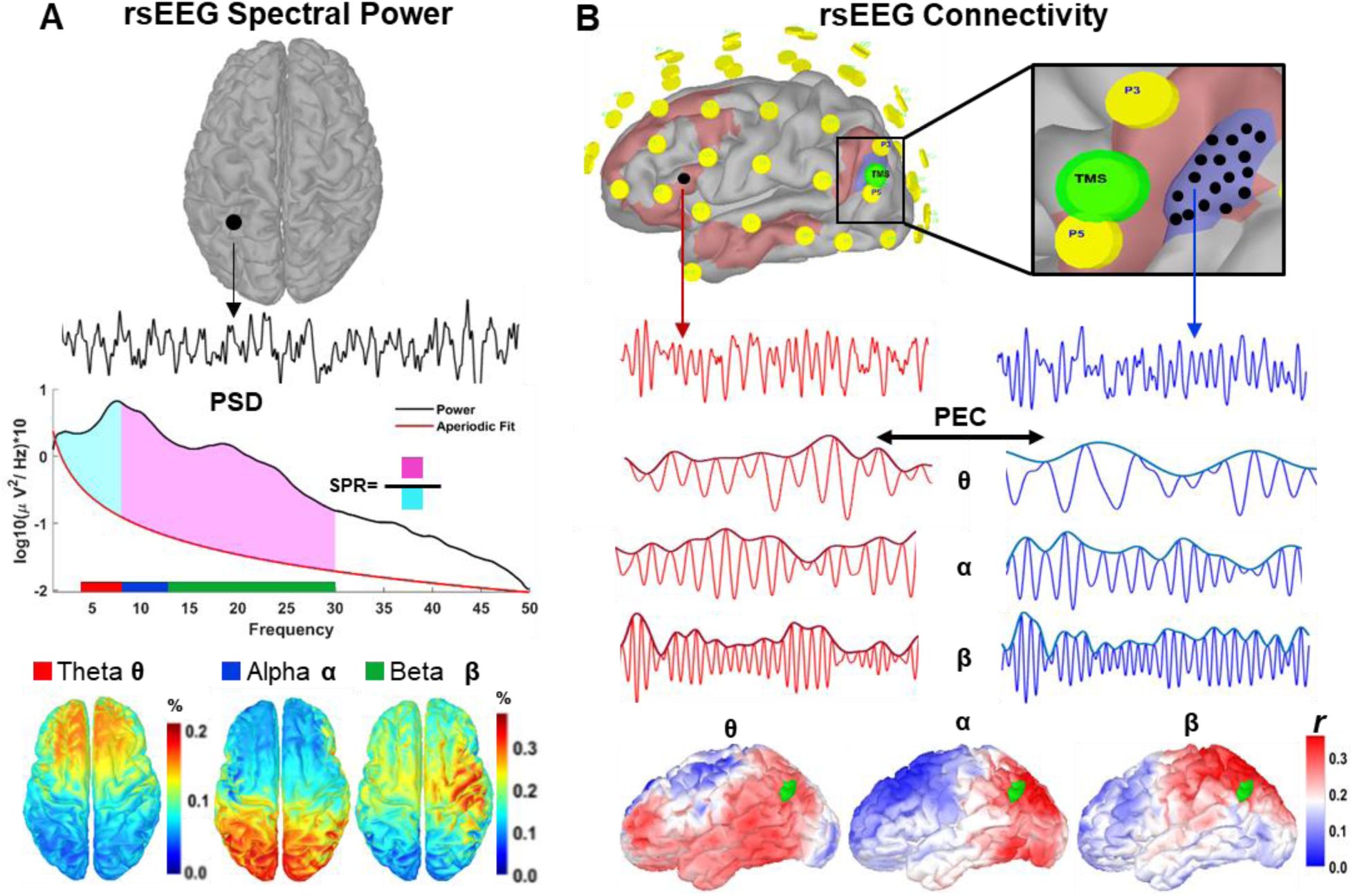
RsEEG Measures. **A:** Current density time series (upper panel) were extracted from each vertex (n=15000) to compute PSD from 1-to-50 Hz (middle panel). PSD exponent (red line in middle panel) is computed and removed from PSD (black line in middle panel) and relative power at canonical EEG frequency bands were calculated at each vertex on the individual cortical surface (lower panel). SPR is calculated as the ratio of delta + theta power (cyan shaded region in middle panel) to the alpha + beta power (magenta shaded region in middle panel) for each vertex. **B:** TMS coordinates (green circle in upper panels) for Left-IPL were used to determine seed vertex on the individual cortical surface. The seed vertex is expanded over the cortical surface to define a seed region (blue shaded region in the expanded upper panel). PEC were computed between each vertex within the seed region (black dots within the blue shaded region) and rest of the brain to generate individual connectivity matrices at delta-theta (2-7 Hz), alpha (8-12 Hz) and beta (13-29 Hz) bands (middle panels with color coded EEG time series). Connectivity matrices were averaged and mapped on individual cortical surface to estimate rsEEG connectivity of the IPL with the rest of the brain (lower panels showing cortical connectivity maps).

### Resting state based neurophysiological abnormalities in AD are not network specific

We performed source reconstruction of rsEEG data (Fig. 5) and computed spectral power density from 1-50 Hz at each vertex across the cortex, defining neural slowing as the spectral power ratio (SPR) between alpha-beta and delta-theta frequencies (Fig. 5A). We also estimated vertex-wise neural synchrony between the TMS site at IPL and the rest of the brain within Delta-Theta (1-7 Hz), Alpha (8-12 Hz), and Beta (13-29 Hz) frequency bands by calculating power envelope correlations (PEC) in individual source space (Fig. 5B). To compare our rsEEG measures of neural synchrony with TMS-EEG measures of network connectivity, we focused on the rsEEG connectivity profile of the left IPL with the rest of the brain (Fig. 5B).

We computed SPR differences between HC and AD groups across entire cortical surface (global) and within local masks of IPL, M1 and frontal DMN (Fig. 6A). We found reduced SPR in the AD group in global (*t*_(79)_=-2.36, *p*=0.020, *d*=0.53), IPL (*t*_(79)_=-2.20, *p*=0.030, *d*=0.49) and M1 (*t*_(79)_=-2.51, *p*=0.014, *d*=0.56). Vertex-wise comparisons of spectral power and SPR from the template cortical surface showed increased delta power over visual regions and increased theta power widespread across the cortex in the AD group. AD participants demonstrated reduced alpha power over temporal and frontotemporal regions, reduced beta power over most of the cortex, and reduced gamma power primarily over lateral-frontal cortices. SPR comparisons revealed significant differences over temporal-parietal regions in the right hemisphere and lateral-frontal regions in the left hemisphere (Fig. 6B).

**Fig. 6.**
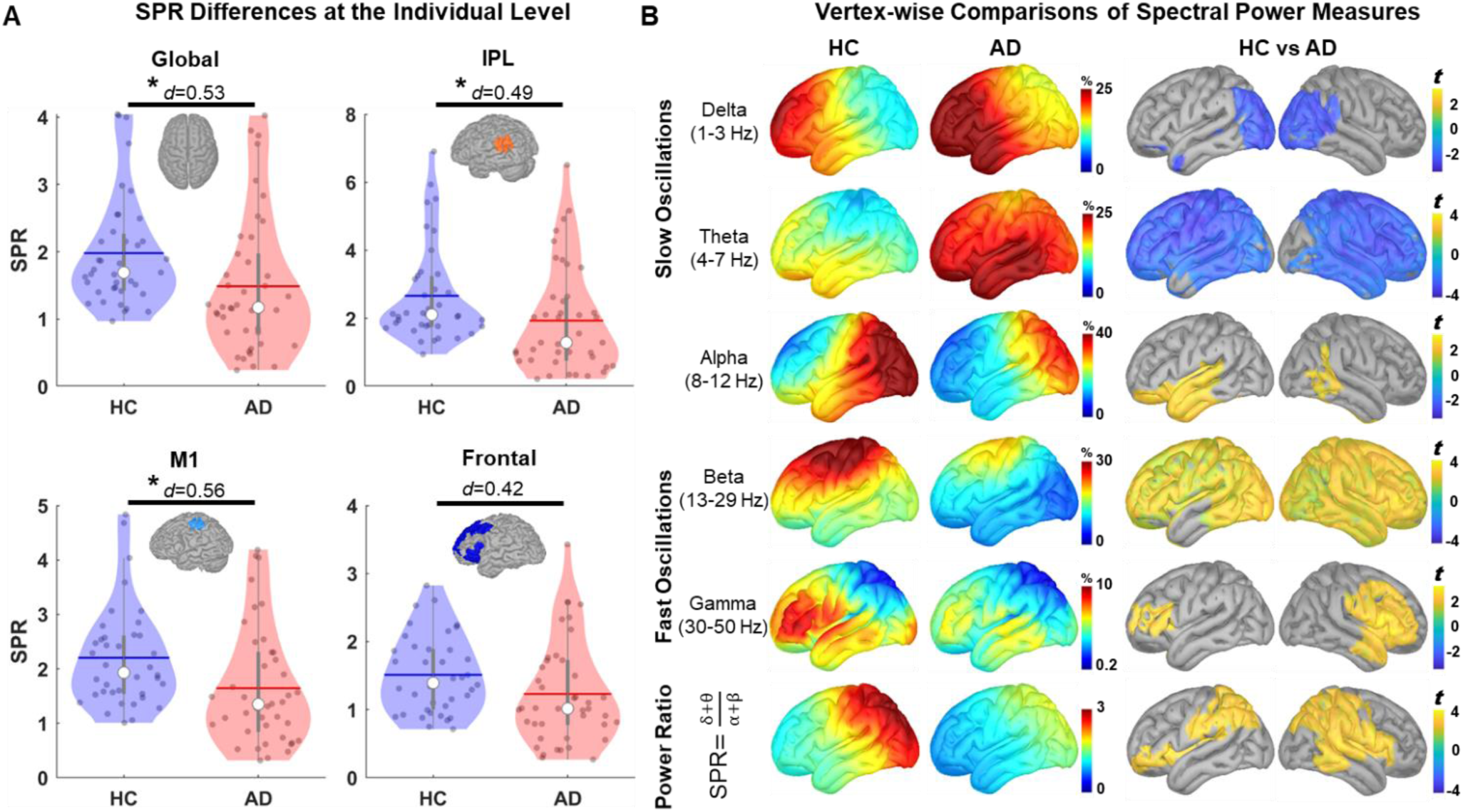
Global slowing in resting state neural oscillations in AD. **A:**Cortical maps showing vertex-wise distribution of spectral power in canonical frequency bands and (left upper panels) and SPR (lower left panels). Lower panels show statistical results of thresholded cluster-based permutation t-tests (cluster *p*<0.05) withhot colors indicating AD < HC and cold colors in indicating AD > HC.**B:** Violin plots showing spectral power ratio averaged across the entire cortical space (global), within individualized masks of IPL, M1 and frontal node of the DMN both in healthy (blue)and AD (red) participants. * in upper violin plot denotes statistical significance at *p*<0.05 with corresponding effect size calculated using Cohens’ *d*.

Neural synchrony analyses (Fig. 7A) showed significantly reduced alpha synchrony across all cortical sites in the AD group (Global: t(79)=-2.70, *p*=0.008, *d*=0.62; IPL: t(79)=-3.03, *p*=0.003, *d*=0.68; Frontal: t(79)=-2.53, *p*=0.013, *d*=0.59; M1: t(79)=-2.72, p=0.008, d=0.62). Similarly, beta band neural synchrony was reduced across all comparison sites (Supplementary Fig. 7), indicating a global decrease in synchrony with no network- or frequency-specific group differences. Group level vertex-wise analyses showed reduced alpha and beta band synchrony primarily over visual, parietal, temporal, and frontal regions (Fig. 7B).

**Fig. 7.**
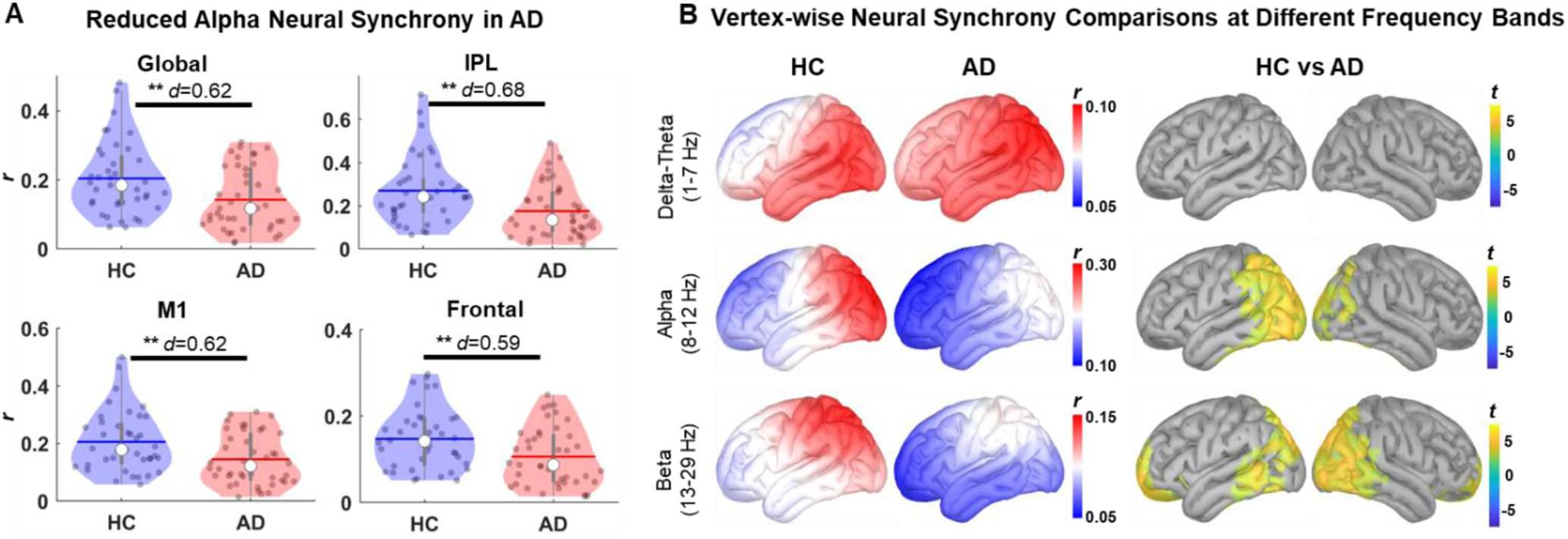
Resting-state neural synchronization differences between AD and HC participants are not network specific. **A:** Violin plots showing alpha band neural synchrony averaged across the entire cortical space (global), within individualized masks of IPL, M1 and frontal node of the DMN both in healthy (blue) and AD (red) participants. * in upper violin plot denotes statistical significance at *p*<0.05 with corresponding effect size calculated using Cohens’ *d*. **B:** Cortical maps showing vertex-wise distribution of neural synchrony in canonical frequency bands and (left panels). Right panels show statistical results of thresholded cluster-based permutation t-tests (cluster *p*<0.05) with hot colors indicating AD < HC and cold colors in indicating AD > HC.

### Global deficits in oscillatory neural activity and neural synchrony correlate to poor cognition in AD

We first examined relationships between potential confounding variables of ‘age’, ‘education’ and ‘cortical thickness’ with RsEEG measures of SPR and neural synchrony in the alpha band (Supplementary Table. 3). IPL cortical thickness was significantly related to increased SPR within the IPL-DMN (*r*=0.39, *p*=0.045). No other significant correlations were found for age and education.

SPR within the IPL-DMN was moderate correlate of better global cognition (ADAS-Cog: *r*=-0.41, *p*=0.006), verbal memory (RAVLT-Total: *r*=0.43, *p*=0.004) and semantic memory (Fluency: *r*=0.38, *p*=0.011), but was not related to working memory function (Fig.8A). Controlling for age and cortical thickness, however, hierarchical regression analyses showed that SPR within the IPL-DMN was not a significant predictor of any cognitive function (*p* > 0.5), explaining an additional 3%, 9% and 3% of the unique variance in ADAS-Cog, RAVLT-Total and Fluency scores, respectively. For the correlations between neural synchrony and cognition, we found that increased alpha synchrony within IPL-DMN is a weak correlate of better global cognition and memory with no significant relationships to any cognitive function (Fig.8A, right panels). Increased beta synchrony in the IPL is a moderate correlate of better global and memory functions, and weak correlate of working memory in the AD (Supplementary Fig.8). Finally, we examined network specificity of significant correlations between IPL SPR and cognition and found almost identical correlation patterns with cognition using global SPR or SRP from the frontal-DMN (black and blue colored scatters in Fig.8A and Supplementary Fig. 8), suggesting that rsEEG measures may be a global signature of decline in global cognitive and memory functions with no distinct network specific relationships. Our vertex-wise correlations support such conclusions by showing a uniform correlation pattern between SPR and cognitive functions across the cortical surface (Fig.8B).

**Fig. 8.**
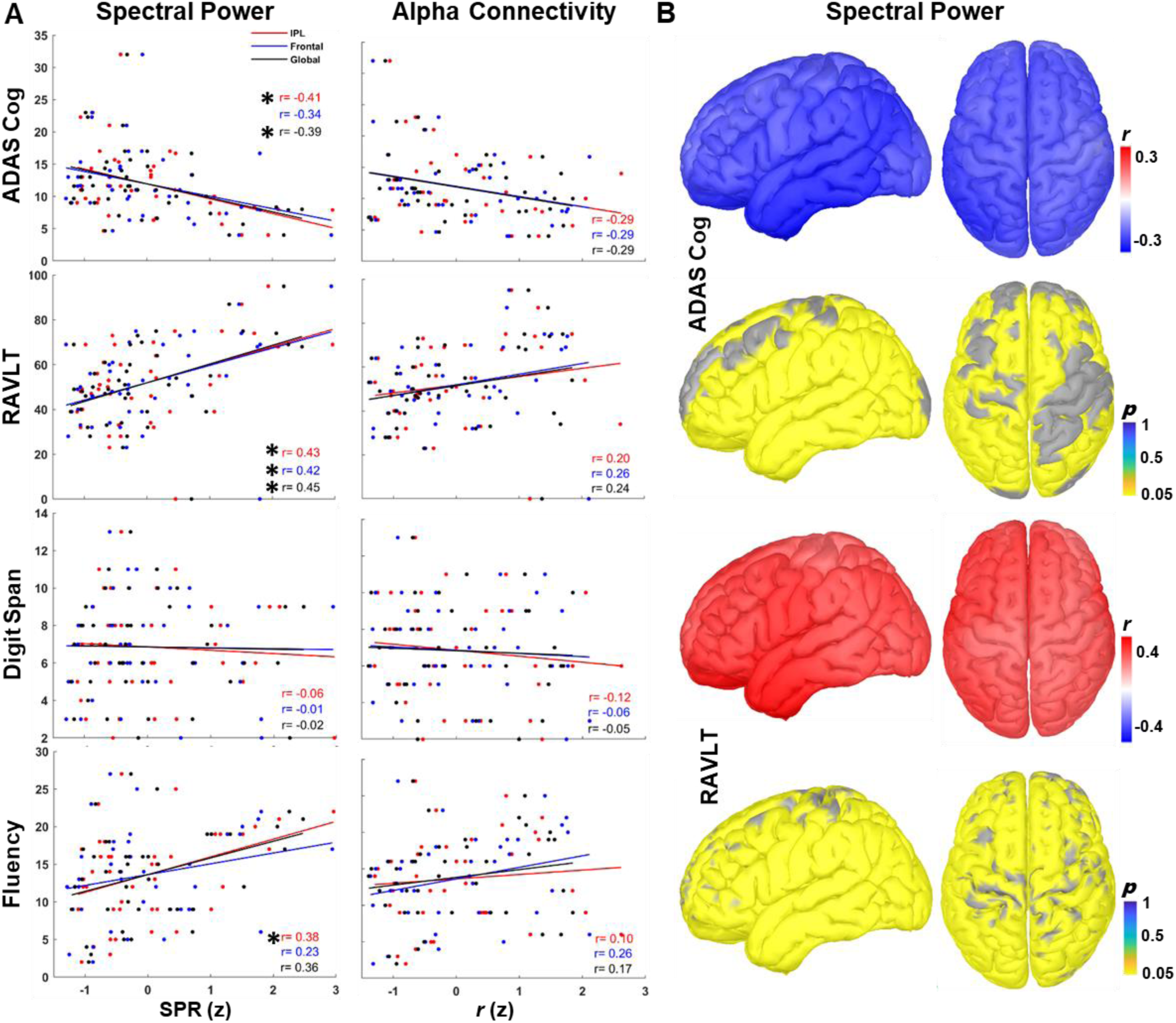
Relationships between resting-state EEG measures of spectral power ratio, neural synchrony and cognitive functions in AD participants. **A:** Scatter plots with regression lines showing bivariate correlations between rsEEG measures of spectral power ratio (left panels) and neural synchrony (right panels) with cognitive function in AD participants. Color codes refer different cortical regions with red showing rsEEG measures from IPL mask and blue showing from the frontal node of the DMN and black showing the average measure computed across the entire cortex (global). Correlation coefficients are provided for each regression line with asterisks indicating statistically significant correlations (*p*<0.05). **B:** Cortical maps showing vertex-wise correlations of SPR with ADAS-cog (upper panels) and RAVLT (lower panels) scores with hot colors indicating positive and cold colors indicating negative correlations. Statistically thresholded cortical maps (*p*<0.05) shows brain regions with significant correlation

## Discussion

Increasing evidence suggests that Alzheimer’s disease (AD) pathology is associated with both cortical hypeexcitability and functional brain network failure, particularly involving early aberrant changes within the DMN^1,16,60–64^. Despite extensive research, our understanding of DMN dysfunction in AD has largely been limited by the constraints of resting-state functional neuroimaging. Here, we developed a novel perturbation-based TMS-EEG approach to characterize cortical excitability with high spatial precision at the site of stimulation and to assess causal connectivity dynamics reflecting the integrity of multi-synaptic signaling within the DMN. We also assessed spontaneous brain activity using commonly applied rsEEG spectral and connectivity measures to comprehensively examine the neurophysiology of AD in relation to cognitive function and structural brain integrity. We found that source localization of TMS-evoked EEG responses revealed local hyper-excitability at the stimulated parietal node with increased propagation to the ipsilateral mesial temporal lobe, along with a breakdown of connectivity between parietal and frontal-DMN regions in AD participants. Most importantly, these TMS-EEG measures were specific to DMN stimulation, served as independent predictors of distinct cognitive functions with spatial and temporal specificity, explain a substantial portion of the variance in individual cognitive function, and may mediate the link between structural brain integrity and cognitive decline in AD. In contrast, although spontaneous neural slowing and disrupted neural synchrony were clearly evident in AD participants, these rsEEG measures and their relationships to cognition were not specific to the DMN but instead reflected global changes in brain neurophysiology. Overall, our results translate neurophysiological signatures of AD observed in animal studies to humans and indicate that perturbation-based measures are sensitive to AD-related alterations in functional brain networks and may serve as biomarkers of distinct cognitive symptoms.

Neurophysiological abnormalities in AD are characterized by a loss of synaptic density and disrupted synaptic transmission^65^, leading to decreased neuronal activity in AD^66^. In mouse models of AD, Palop and colleagues (2007) reported seizure activity in cortical and hippocampal neurons indicating an excitatory effects of Aβ on neuronal function^4^. Subsequent research confirmed these findings, showing clusters of abnormally hyperactive cortical neurons exclusively near Aβ plaques^5^ and in hippocampal neurons even before the formation of plaques, often driven by soluble Aβ oligomers, establishing neuronal hyper-excitability as a key neurophysiological signature in the AD spectrum^67^. However, the presence and relevance of cortical excitability in human AD, particularly within brain networks involved in memory processes, have remained poorly understood. While cortical hyper-excitability in the form of inter-ictal epileptiform discharges have been identified in 20-45% of patients with early AD, where they have been associated with faster declines in global cognition^68,69^, the majority of AD patients lack these findings despite extended monitoring, and network-specific associations with cognition are difficult to establish. In our study, we leveraged the high spatial focality and causal nature of TMS combined with the precise temporal resolution of EEG, alongside advanced simulation and inverse algorithms, to map perturbation-based neurophysiology in the DMN and its links to cognitive functions and brain structure in AD patients. We found that AD participants exhibited increased cortical excitability at the IPL-DMN up to 65ms post-TMS, which extended to left temporal DMN from 60-85ms. Conversely, starting at 75ms, frontal DMN activity was markedly reduced in AD participants compared to controls, suggesting a pattern of hyper-excitable posterior and temporal-DMN with disrupted connectivity to frontal DMN. These findings build on existing evidence showing that parietal and temporal DMN regions are particularly vulnerable to early AD pathology^13,70^. Recent neuroimaging studies have linked these areas to reduced metabolism^14,61^, cortical atrophy^71^, and aberrant DMN connectivity, including hypo-connectivity within medial parietal nodes and hyper-connectivity between medial and lateral temporal-parietal nodes^16,72^. Our results not only translate these imaging-based findings into the cortical neurophysiology of DMN dysfunction but also highlight a novel finding of disrupted posterior-frontal DMN connectivity in AD. The utility of TMS in detecting cortical excitability and connectivity impairments has been well established in patients with stroke^73,74^ and spinal cord injury^75–77^, where motor evoked potentials (MEPs) reflect decreased excitability in the affected hemisphere in stroke, and indicate disrupted signal transmission in spinal cord injury indexing the integrity of the corticospinal tract. Our findings extend this utility to cortical neurophysiology in AD, demonstrating that TMS-EEG can reveal hyper-excitability and disrupted connectivity within the DMN.

Our control analyses using M1- and Sham-TMS data showed no significant differences between the groups, confirming that the abnormally increased local cortical excitability is specific to DMN stimulation and cannot be attributed to non-transcranial effects of TMS. While the lack of significant group differences in motor cortex maps following M1-TMS contrasts with much of the previous literature showing increased cortical excitability in the motor cortex using MEPs^78^, we argue that our local cortical excitability results reflect the severity of network-specific neurophysiological abnormalities in AD. Our supplementary analyses revealed that resting motor threshold (RMT) is significantly lower in AD participants, resulting in weaker E-field strength for both M1 and IPL stimulation (Supplementary Fig. 12). This aligns with a recent meta-analysis identifying RMT as the most robust measure consistently lower in AD participants^78^, whereas findings related to scalp measures of TMS-EEG responses were highly inconsistent^79^. To our knowledge, only one other study has used source localization of TMS-EEG to describe cortical excitability and connectivity in the motor cortex in AD, but this study provided source results only for visualization purposes without statistical analyses^46^. Consistent with our results, TMS of M1 in healthy subjects typically shows spatially distributed and temporally distinct network activation patterns, propagating from premotor to motor and somatosensory cortices, reflecting the anatomical connectivity of the motor cortex. In contrast, AD participants exhibited stronger, longer-lasting local activations that remained mostly within the stimulated area, with poor propagation within the stimulated network^46^. The fact that spTMS elicited statistically similar neural responses at M1 but significantly higher activations at IPL, despite weaker E-field strength and lower stimulation intensities in AD participants, reflects the network-specific severity of neuropathological alterations across the spectrum of disease progression. Given that AD pathology first manifests in the parietal brain regions^17^ and spreads to other cortical regions^6,72^, with the DMN being one of the earliest and most affected networks, our findings of increased IPL and temporal DMN excitability in early AD participants align with this temporal progression. The significantly altered cortical excitability and connectivity in DMN regions, but less pronounced abnormalities within the motor cortex, highlight the sensitivity of TMS-EEG in detecting early network-specific changes in cortical excitability and connectivity.

Our analyses of TMS-EEG measures of cortical excitability and network connectivity in relation to specific cognitive functions revealed several key findings. First, we observed strong correlations between increased IPL excitability and global cognitive function, as measured by ADAS-Cog, as well as episodic memory impairment, as measured by RAVLT. Conversely, connectivity between the IPL and frontal DMN nodes was associated most strongly with working memory and executive functions, consistent with the role of the dorsolateral prefrontal cortex (DLPFC) in these cognitive processes^80^. These distinct correlations suggest that IPL excitability is particularly sensitive to global cognitive and memory deficits, while frontal DMN connectivity is more closely related to working memory and executive functions, demonstrating how different TMS-EEG measures are specifically associated with different cognitive domains. Second, we found that early responses (15-65ms) following IPL stimulation, where hyper-excitability was present, were significantly related to global cognition and memory. In contrast, only late responses in the frontal-DMN node were associated with working memory and fluency. This temporal specificity suggests that local excitatory activity, reflecting the immediate neural effects of TMS, links localized disruptions in cortical excitability to global cognition and memory, both of which are closely associated with parietal lobe functions^81^. Later frontal DMN connectivity responses, representing more integrative, trans-synaptic processing, likely reflect the integrity of long-range network connectivity and better index cognitive processes relying on the coordinated activity of frontal and parietal regions, such as working memory^82,83^. Third, the absence of significant correlations with M1 stimulation and sham-TMS demonstrates the specificity of these findings to DMN-related alterations in AD. Importantly, hierarchical regression analyses indicated that IPL hyper-excitability is an independent predictor of global cognitive decline, explaining a substantial portion of the variance in ADAS-Cog after controlling for cortical thickness, age, and education. Cortical thinning in regions such as the medial temporal lobe, lateral parietal cortex, and frontal areas is a well-documented hallmark of AD^71,84,85^, strongly associated with AD pathology and severity^86^. Our findings align with this literature, showing significantly reduced cortical thickness in AD participants (Supplementary Fig. 12), which was also associated with poorer cognitive performance (Supplementary Table 1). However, when cortical hyper-excitability was considered, the relationship between cortical thickness and global cognition was no longer significant, suggesting that cortical hyper-excitability maybe an underlying neurophysiological abnormality that links impairments in structural brain integrity to cognitive decline. Moreover, the unique variance explained by frontal-DMN connectivity in working memory and verbal fluency highlights the importance of intact network communication for preserving these cognitive domains. Overall, these results emphasize the relevance of both temporal and spatial specificity in TMS-EEG measures, highlighting their potential as sensitive biomarkers for AD-related neurophysiological alterations and cognitive decline.

In our study, we also assessed commonly used rsEEG measures to comprehensively characterize the neurophysiological signatures of AD. We observed a global shift in spectral power dynamics, characterized by a decrease in faster alpha-beta oscillations relative to slower delta-theta frequencies, a phenomenon known as ‘neural slowing^87^’. This neural slowing was evident across the cortex, including regions such as the IPL, M1, and frontal DMN. Additionally, we found decreased neural synchrony within the alpha and beta frequency bands, particularly in the visual, parietal, and temporal-frontal regions, indicating widespread disruption in synchronized neural activity in AD. Consistent with these neurophysiological alterations, our correlation analyses revealed that lower SPR and reduced alpha synchrony were moderately associated with global cognitive decline and memory impairments, although these relationships were not network-specific. These findings align with previous studies that have reported similar patterns of neural slowing in AD^20,25^ and have linked disrupted alpha and delta-theta synchrony to AD pathology and cognitive decline^27,88^. However, the global nature of rsEEG measures, which reflect widespread changes in brain physiology, may explain why these metrics fail to capture localized alterations critical for understanding functional network failures in AD. RsEEG is sensitive to diffuse, spontaneous neural oscillations arising from multiple overlapping networks, making it less effective in isolating dysfunctions within specific networks like the DMN. Moreover, the correlational nature of spontaneous oscillations does not directly index key neurophysiological signatures of AD, such as cortical hyper-excitability and causal network dysconnectivity. In contrast, our TMS-EEG approach, which allows for the targeted stimulation of specific brain networks, revealed localized alterations in cortical excitability and captured abnormal causal connectivity dynamics within the DMN. These TMS-EEG measures also demonstrated distinct correlation patterns with different cognitive functions, suggesting that perturbation-based neurophysiological modalities provide a more precise characterization of network-specific dysfunctions in AD, offering deeper insights into how these dysfunctions relate to specific cognitive deficits.

While our findings provide important insights into the neurophysiological alterations associated with AD, several limitations should be acknowledged. A key limitation is our inability to quantitatively relate our findings to pathological markers such as regional Aβ and p-tau levels. Integrating these biomarkers could have offered a more comprehensive understanding of the neurobiological mechanisms driving the observed changes in cortical excitability and connectivity. Additionally, our study did not differentiate between AD subtypes, such as typical amnestic AD, primary progressive aphasia, and posterior cortical atrophy, each of which exhibits distinct patterns of cognitive impairment and neurodegeneration. This limits our ability to generalize our TMS-EEG findings across the spectrum of AD presentations and suggests a need for future research to explore whether TMS-EEG measures can differentiate between these subtypes. The cross-sectional design of our study also restricts our ability to determine whether TMS-EEG measures are sensitive to tracking disease progression or responses to treatment over time. Longitudinal studies are necessary to validate the utility of these measures as biomarkers for monitoring disease progression and therapeutic efficacy. Despite these limitations, the robust correlations we observed between TMS-EEG measures and cognitive function, which exceeded those typically reported in fMRI^16^ and MRI studies, suggest that TMS-EEG may offer a more sensitive approach than resting state neurophysiology for detecting network-specific dysfunctions in AD, even with a limited sample size.

Taken together, our findings demonstrate that perturbation-based measures are powerful tools for characterizing network-specific neurophysiological alterations in AD, offering insights that extend beyond traditional resting-state and structural measures. By highlighting the role of cortical hyper-excitability and disrupted connectivity within the DMN, our study provides a deeper understanding of the functional network failures underlying cognitive decline in AD. Future research should focus on integrating these neurophysiological measures with pathological markers and assessing their utility in tracking disease progression across different AD subtypes in longitudinal cohorts. This approach could refine our understanding of AD pathophysiology and potentially guide the development of targeted interventions.

## Methods

### Participants

Forty-two biomarker-positive early AD (18 female, Age_mean_=70.95± 7.63 years) and 40 healthy older adult control (HC) participants (19 female, Age_mean_=70.63 ± 6.41 years) were included in this study. Inclusion criteria for early AD participants were: aged 50-90 years old, on a stable dose of medications for memory loss (e.g. donepezil, rivastigmine or memantine) as defined by 4 consecutive weeks of treatment at a fixed dose, meeting the NINCDS-ADRDA criteria for probable AD, Mini Mental State Examination (MMSE) ≥ 20 (to insure ability to give appropriate consent), positive AD biomarker status (as defined by CSF biomarkers or florbetapir amyloid PET study) and clinician dementia rating (CDR) of 0.5-1.0. Inclusion criteria for HC participants were: aged 50-80 years old, Normal neurologic exam, MMSE > 28 and CDR of 0-0.5 (CDR of 0.5 was allowed if the participant was found to have only subjective memory complaints by consensus discussion between the study MD and neuropsychologist. Subjective memory complaints were defined as there being no objective evidence of cognitive impairment or memory loss on cognitive testing or informant report. In this sample, only one HC participant had CDR of 0.5, Supplementary Table. 1). Exclusion criteria included a diagnosis of epilepsy, current or past history of any neurological disorder (other than dementia in the AD group); stroke; intracranial brain lesions; history of previous neurosurgery; head trauma that resulted in residual neurologic impairment; or substance use disorders within the past six months. An abnormal neurologic or cognitive exam and use of medications that could alter cortical excitability, as determined by the investigators, were additional exclusion criteria for HC participants only. Experimental protocols and voluntary participation procedures were explained to all participants before they gave their written informed consent to the study which conformed to the Declaration of Helsinki, and had been approved by the Institutional Review Board of the Beth Israel Deaconess Medical Center. If an AD participant was interested in participating in the study but was determined to be unable to consent by the study MD, a legally authorized representative provided informed consent. RsEEG data were collected from all participants. TMS-EEG data were available in 37 AD participants. RsEEG data were collected from all participants. TMS-EEG data were available in 37 AD participants.

### Data acquisition

A T1-weighted (T1w) anatomical MRI scan was obtained for all participants and used for neuronavigation. MRI data were acquired on a 3T scanner (GE Healthcare, Ltd.) using a three-dimensional spoiled gradient echo sequence: 166 axial-oriented slices for whole-brain coverage; 240-mm isotropic field-of-view (FOV); 0.937-mm × 0.937-mm × 1-mm native resolution; flip angle = 15°; echo time (TE)/repetition time (TR) ≥ 2.9/6.9 ms; duration ≥432 s.

TMS was delivered using a figure-of-eight−shaped coil with dynamic fluid cooling (MagPro 75-mm Cool B-65; MagVenture A/S) attached to a MagPro X-100 stimulator (MagVenture A/S). For single-pulse sham TMS stimulation, a MagVenture Cool-B65 A/P coil was used with a 3 cm thick plastic spacer. Individual high-resolution T1w images were imported into the Brainsight TMS Frameless Navigation system (Rogue Research Inc.), and coregistered to digitized anatomical landmarks for online monitoring of coil positioning. Whole-scalp 64-channel EEG data were collected both for rsEEG and TMS-EEG recordings at a sampling rate of 5000 Hz with a high frequency cut-off at 1350 Hz using a TMS-compatible amplifier system (actiCHamp system; Brain Products GmbH) and labeled in accordance with the extended 10–20 International System. EEG data were online-referenced to AFz electrode. Electrode impedances were maintained below 5 kΩ. EEG signals were digitized using a Brainamp actiCHamp amplifier and linked to BrainVision Recorder software (version 1.21) for online monitoring. Digitized EEG electrode locations on the scalp were also coregistered to individual MRI scans using Brainsight TMS Frameless Navigation system. Motor-evoked potentials (MEPs) were recorded from the right first dorsal interosseous (FDI) and the abductor pollicis brevis (APB) muscles following single pulse TMS of left motor cortex. Ag−AgCl surface electrode pairs were placed on the belly and tendon of the muscles and a ground was placed on the right ulnar styloid process.

### TMS Targets

We used group-level functional parcellations and confidence maps on the Montreal Neurological Institute (MNI) template brain to target the most consistent regions within the angular gyrus (Left-<u>IPL</u>: −55.1, −70.5, 27.7) that had the highest likelihood of occurring in the DMN. A custom processing pipeline was developed to take each subject’s anatomical MRI, create a non-linear transform from subject to MNI space and then use the inverse of that transform to bring the coordinates back into subject space using FSL’s “FNIRT” tool. The transformed coordinates along with individual high-resolution T1w images were then imported into the Brainsight™ TMS Frameless Navigation system (Rogue Research Inc., Montreal, Canada), and co-registered to digitized anatomical landmarks for online monitoring of coil positioning. Left motor cortex (L-M1) was stimulated at the TMS motor hotspots for each individual as described in detail below.

### Experimental Procedures

Throughout the session, participants were comfortably seated in an adjustable chair. At the beginning of the TMS visit rsEEG recordings were performed first in eyes closed (EC) condition for five minutes. Participants were instructed to remain awake throughout this period, and were intermittently instructed to blink several times to help maintain alertness. Following rsEEG recordings, the motor hotspot for eliciting MEPs in the right FDI muscle was determined by delivering single TMS pulses and moving the TMS coil in small incremental steps after two to three stimulations in each spot, over the hand region of left motor cortex with 45° rotation in relation to the parasagittal plane (inducing posterior-to-anterior current in the underlying cortex). The hotspot was defined as the region where single-pulse TMS elicits larger and more consistent MEPs in the FDI muscle, as compared to the APB muscle, with the minimum stimulation intensity. The FDI hotspot was digitized on each participant’s MRI image. Resting motor threshold (RMT) was determined on the FDI hotspot as the minimum stimulation intensity eliciting at least five MEPs (≥50 μV) out of 10 pulses in the relaxed FDI using biphasic current waveforms. In compliance with the International Federation of Clinical Neurophysiology safety recommendations^89^, participants were asked to wear earplugs during hotspot and RMT trials to protect their hearing, and to minimize external noise. TMS was administered with a thin layer of foam placed under the coil to minimize somatosensory contamination of the TMS-evoked EEG potentials. To minimize AEPs related to the TMS click, auditory white noise masking was used throughout the TMS stimulation. The intensity of noise masking is determined as the highest noise level participants could tolerate below 90db. Following determination of RMT, a total of 150 single TMS pulses were delivered to each stimulation target (DMN node target in the left inferior parietal lobule IPL-TMS and motor hot spot in the left motor cortex M1-TMS) at an intensity of 120% RMT with randomly jittered (3,000 to 5,000ms) interstimulus intervals. Sham-TMS was administered on the motor hot spot of the FDI muscle over left M1. An active/sham TMS coil (Cool-B65 A/P, MagVenture A/S, Farum, Denmark) was flipped to the placebo side and stimulation intensity was kept identical to actual TMS, but with induced currents on the opposite vertical direction to the targeted gyri (Supplementary Fig. 9). A 3D printed 3 cm spacer was attached to the placebo side (MagVenture A/S, Farum, Denmark) of the coil to further ensure the elimination of residual currents on the placebo side of the coil. White noise masking was presented through earplug-earbuds at the maximum volume comfortable for each participant. Small current pulses between 2 and 4 mA and proportional to the intensity of actual TMS pulse were delivered over the left forehead, over the frontalis muscle, using surface electrodes (Ambu Neuroline 715 12/Pouch) to approximate somatosensory sensations arising from skin mechanoreceptors and scalp muscles during active-TMS condition (Supplementary Fig.9). Operators continuously monitored participants during the TMS blocks for their wakefulness, and prompted them to keep their eyes open and stay fully relaxed in case of visible signs of drowsiness or tension.

### EEG preprocessing

All EEG data pre-processing was performed offline using EEGLAB 22.1^90^, and customized scripts running in Matlab R2022b (Math-Works Inc., USA). rsEEG recordings were first low-pass filtered at 100Hz using a forward-backward 4^th^ order Butterworth filter and down sampled to 250 Hz. Visual inspection was performed to identify and reject extremely noisy channels (1.88 ± 1.36 channels were deleted on average; range 0–5 out of 63). rsEEG data were then segmented into 3000 ms epochs and each epoch was tagged based on voltage (≥100 mV), <u>kurtosis</u> (≥3), and joint probability (Single channel based threshold ≥3.5sd; All channel-based threshold ≥ 5sd) metrics to identify excessively noisy epochs. Visual inspection was performed on the tagged epochs for the final decision for the removal of noisy epochs (6.75 ± 8.23 epochs were deleted on average). RsEEG data were then notch filtered between 57 and 63Hz, band pass filtered using a forward-backward 4^th^ order Butterworth filter from 1 to 100Hz, and referenced to global average. To avoid edge effects of the Butterworth filter at individual epochs, each epoch was first replicated, inverted and then reflected back to both ends of the signal for symmetrical extension of the original epoch before filtering. Original epochs were then extracted back following the filtering process for further processing steps. A fast <u>independent component</u> <u>analysis</u> (fICA) was performed to manually remove all remaining artifact components including eye movement/blink, muscle noise (EMG), single electrode noise, and cardiac beats (EKG) (see Supplementary Fig. 10 for details of identifying ICA components). A semi-automated artifact detection algorithm “tesa_compselect” incorporated into the open source TMS-EEG Signal Analyzer TESA v0.1.0-beta extension^91^ for EEGLAB was used to visually inspect and manually remove artifactual resting-state EEG ICA components based on their frequency, activity power spectrum, amplitude, scalp topography, and time course (http://nigelrogasch.github.io/TESA/). In average, 19.15 ± 7.02 out of 63 ICA components were removed across all data sets and visits. Rejected channels were interpolated using spherical interpolation. rsEEG data from one HC participant were discarded from the analyses because of extreme EMG contamination across most of the channels. Final sample size for rsEEG analyses consists of 42 AD and 39 HC participants.

### TMS-EEG preprocessing

TMS-EEG data were segmented into 3000 ms epochs, each starting 1000 ms before (pre-stimulus) and ending 2000 ms (post-stimulus) following TMS pulse, respectively. Baseline correction was performed by subtracting the mean pre-stimulus (−500 to −100) signal amplitude from the rest of the epoch in each channel. Following baseline correction, data were visually inspected to identify noisy channels (2.26 ± 1.38 channels were deleted on average; range 0–5 out of 63). Zero-padding between −2 ms and 10 ms time range was then applied to remove the early TMS pulse artifact from the EEG data. All zero padded epochs were then tagged based on voltage (≥100 μV), <u>kurtosis</u> (≥3), and joint probability (Single channel-based threshold ≥ 3.5sd; All channel-based threshold ≥ 5sd) metrics to identify excessively noisy epochs. Visual inspection was performed on the tagged epochs for the final decision for the removal of noisy epochs (7.89 ± 8.59 epochs were deleted on average). Next, an initial round of fICA was performed using TESA v0.1.0-beta extension^91^ to identify and remove components with early TMS evoked high amplitude electrode and EMG artifacts (1 ± 1 components were removed; range 0–3 out of 63). After the first round of fICA, the EEG data were interpolated for previously zero-padded time window around TMS pulse (−2 ms–10 ms) using linear interpolation, band pass filtered using a forward-backward 4th order butterworth filter from 1 to 100 Hz, notch filtered between 57 and 63 Hz, and referenced to global average. Missing/removed channels were interpolated using spherical interpolation, and sham stimulation blocks were merged with IPL and M1 blocks separately. Subsequently, a second round of fICA was run to manually remove all remaining artifact components including eye movement/blink, muscle noise (EMG), single electrode noise, TMS evoked muscle, cardiac beats (EKG). We performed a sham-informed ICA-based process for identifying and removing auditory evoked potentials in IPL and M1 data sets as previously described^92^. Briefly, ICA components that met the following criteria were classified as AEP components and removed. (1) The time-course has three peaks, P50-N100-P200, with the lowest amplitude in the P50 compared with N100 and P200; (2) the scalp topography reflects left/right symmetry and a central distribution anterior to Cz; and (3) the component is shared across both active stimulation sites and sham stimulation. Details for identifying and validating AEP components are provided in Supplementary Fig. 11. TMS-EEG data from four AD participants and 1 HC participant were discarded from the analyses due to poor data quality. Final sample size for TMS-EEG analyses consists of 33 AD and 39 HC participants.

### EEG source reconstruction

Both rsEEG and TMS-EEG source reconstruction was performed using Brainstorm software^93^. First, individual cortical surface reconstruction was performed by running Freesurfer “recon-all” command on T1-weighted anatomical MRI scans. Group-averaged functional cortical atlases (n = 1,000) consisting of seven resting-state networks were also morphed into the subject’s cortical surface using surface-based registration in freesurfer (Fig. 1A). Next, digitized EEG channel locations and anatomical landmarks of each subject were extracted from Brainsight™ (nasion ‘NAS’, left pre-auricular ‘LPA’, and right pre-auricular ‘RPA’ points), and registered onto individual MRI scans in brainstorm. Continuous rsEEG and epoched TMS-EEG, −500 ms–1000 ms with respect to TMS pulse for each TMS trial, were uploaded and forward modeling of neuro electric fields was performed using the open MEEG symmetric boundary element method, all with default parameter settings. For TMS-EEG, noise covariance was estimated from individual trials using the pre TMS (−500 to 0) time window as baseline while the first 10 seconds of rsEEG data was used for noise modelling of rsEEG data. The inverse modeling of the cortical sources was performed using the minimum norm estimation (MNE) method with current density maps for rsEEG and with dynamic <u>statistical parametric</u> <u>mapping</u> (dSPM) for TMS-EEG as the source estimation measures. We choose dSPM for TMS-EEG as it further normalize current density maps to the baseline period (preTMS) with z-scores and provide a statistical map of TMS evoked responses. Source dipoles for both measures were constrained to the individual cortical surface.

### Generation of surface based masks

Currently, electric-field (E-field) modeling of TMS is considered as the gold standard for estimating the extent of local cortical regions directly activated by TMS^94^. Therefore, we performed E-field simulations on individual brain models to identify TMS-activated local cortical regions and generate local surface-based masks for measuring local cortical excitability at the source level (Fig. 1B, left upper and middle panels). First, we performed E-field simulations in SimNIBS software^57^ (version 4.1) using individual brain models from FreeSurfer. We used the ‘MagVenture Cool-B65’ TMS coil model from Drakaki et al., with dI/dt values of the TMS machine set to 120% RMT (actual stimulation intensity) and with an 8 mm skin-to-coil distance to account for the space of EEG electrodes on the scalp. The coil position in SimNIBS was set to the 3D world coordinates of each TMS site on the individual MRI model, derived from Brainsight. AD participants had significantly lower peak E-field strength than HC participants (*t*_(70)_ = −2.68, *p* = 0.009, Supplementary Fig. 12). To better understand significant E-field strength differences between the groups, we compared RMT and scalp-to-cortex distances and found that while RMT was significantly lower in AD participants (*t*_(70)_ = −2.15, *p* = 0.034), scalp-to-cortex distances were statistically similar between the groups (*p* > 0.5), indicating lower RMT as the source of E-field differences between the groups (Supplementary Fig.12B).

After performing SimNIBS simulations, the E-field distribution for each TMS condition (Left-IPL and M1) at individual volume space was projected onto the cortical surface model using FreeSurfer’s^95^ mri_vol2surf function. Following projection, we employed the mri_binarize function to threshold the surface data, retaining only values greater than 0.01% of the maximum value to identify local cortical regions most affected by TMS. Subsequently, we generated surface masks from the binarized data using mri_cor2label, which converted the thresholded surface data into label files representing local anatomical regions. These surface masks were further refined and annotated with FreeSurfer’s mri_label2label and mris_label2annot functions, enabling precise delineation of each local cortical mask. Finally, annotation files for each site were uploaded to Brainstorm with individual brain models from FreeSurfer and used as scouts for assessing local cortical excitability (Fig. 1B, right upper and middle panels). These surface masks were also used to compute average cortical thickness (in mm) within the defined regions using the mris_anatomical_stats function in FreeSurfer, allowing us to quantify the local cortical structure in relation to TMS activation. In line with expectations, AD participants had significant cortical thinning (*t*_(70)_ = −5.20, *p* = 0.000) at the parietal DMN regions (Supplementary Fig. 12B).

### TMS-EEG measures

Using e-field-based local masks and fMRI-connectivity-based DMN masks, we extracted average current density time-series (1500ms) representing a 500ms baseline period and a 1000ms post-TMS activity period. These time series were rectified to obtain absolute values and converted into z-scores by normalizing the source activity at each sample to the pre-TMS period (0-500ms baseline). For assessing local cortical excitability, we utilized z-score EEG activity time series derived from e-field-based local masks. We first performed sample-wise independent t-tests on the z-score EEG time-series to characterize cortical excitability differences between the groups at millisecond temporal resolution (Fig. 2C, left panels). Initially, we focused on identifying significant differences within the first 50ms following TMS, as this window often captures the earliest TEPs at the site of stimulation. If significant sample clusters were detected within the first 50ms, we expanded the analysis to define a broader time window that includes these clusters and extends to encompass the full range where the average EEG activity time series showed visible differences between groups. For example, the dashed black line in Fig. 2C (upper left panel), marks the 15-65ms window, is selected to capture both significant clusters and adjacent time points where EEG activity differed between groups. This broader time window was then used to compute the average z-score EEG activity for further analyses (illustrated by the violin plots on the right panel of Fig. 2C, showing the distribution of average z-score EEG activity within this window across participants). By defining this broader data-driven window for further analysis, we ensure a comprehensive evaluation of local cortical excitability that incorporates both the statistically significant clusters and the broader range of group differences observed in the time-series data. For whole brain analyses, we also used these windows to first compute average activity across the entire surface and project average surface activity to MNI template for each participant. For network connectivity measures, we searched for significant time windows after 50ms following TMS and applied the same procedures described above. We defined network connectivity as the average amount of activity extracted from non-stimulated regions of the DMN, representing how much activity propagated from the stimulated to other nodes of the network (Fig. 1B, lower panel, and Fig. 1C). This metric can be conceptualized as a proxy for network connectivity, with higher values indicating increased connectivity between the stimulated and non-stimulated nodes of the DMN.

### rsEEG measures

For resting-state EEG (rsEEG) power measures, we first estimated the broadband (1-50 Hz) spectral power density (PSD) of current density time series at each vertex (n = 15,000) using the Welch method^96^. We applied a 2000 ms Hamming window with a 50% overlap ratio for the PSD estimations (Fig. 5A). Next, we parameterized the EEG power spectra at each vertex into periodic and aperiodic components using the “Fitting Oscillations One Over F” (FOOOF) toolbox^97^ (Fig. 5A, middle panel). After parameterization, we removed the aperiodic component (exponent) from the spectra to focus solely on the periodic (oscillatory) components. This approach helps prevent misinterpretation of oscillatory power changes that could result from shifts in the aperiodic exponent rather than true oscillatory dynamics. By focusing on the periodic components, we ensure that our analysis reflects actual oscillatory activity across canonical frequency bands without the confounding influence of non-oscillatory background activity (Fig. 5A lower panel). This methodological choice aligns with recent advancements in neural signal analysis, which emphasize the importance of distinguishing between periodic and aperiodic components to better understand neural dynamics and their alterations in clinical populations such as those with Alzheimer’s disease^97–102^. After removing the aperiodic slope (exponent) from EEG power spectra, we computed relative spectral power at canonical frequency bands (Δ-delta: 1-3 Hz, θ-theta: 4-7 Hz, α-alpha: 8-12 Hz, β-beta: 13-29 Hz and γ-gamma: 30-50 Hz) as the percentage (%) of the total absolute power expressed (1-50Hz) at each vertex (Fig. 5A lower panel). Absolute power estimates were used to compute the spectral power ratio (SPR), defined as the ratio of power in α and β to power in Δ and θ: (α+ β)/(Δ+ θ). Lower SPR values indicates the degree of *“neural slowing”*. This approach is particularly useful for assessing alterations in the spectral dynamics of EEG, capturing the global shift in the EEG power spectrum from higher to lower frequencies, a pattern previously reported in AD^20^.

For rsEEG connectivity, we computed seed-based orthogonalized power envelope correlations^103^ (PEC) at individual source space using current density time series (Fig. 2B). The PEC approach has been introduced to measure functional brain connectivity at distinct neural oscillations using MEG/EEG signals, is sensitive for characterizing abnormal brain dynamics in several psychiatric and neurological conditions with correlated changes in cognitive behavior^99,104–106^, and is the most robust and reproducible connectivity metric compared to phase, spectral coherence and auto-regressive based connectivity estimates^106^. To characterize rsEEG connectivity of the stimulated DMN node, we used individual TMS coordinates of Left-IPL to generate connectivity seed regions for each participant. We first found the seed vertex on the individual cortical surface that was closest to the TMS coil over the scalp, and expanded the seed vertex over the cortex in all directions to the closest 40 vertices (roughly 3cm^2^) to define the seed region for each individual (Fig. 5B upper panels). We then iteratively computed PEC as the *“neural synchrony”* measure between each vertex within the seed region and rest of the brain vertices at delta-theta (2-7 Hz), alpha (8-12 Hz) and beta (13-29 Hz) frequencies. Current density time series were narrow-band pass filtered at each frequency band and Hilbert transformed to compute analytical signals as previously described^107^ (Fig. 5B middle panels). Analytical signals, a complex valued time series data containing momentary phase and amplitude information, within each vertex are orthogonalized with respect to the seed vertices. Power envelopes were then extracted from each orthogonalized analytical signal and seed vertices and transformed to natural logarithm to improve normality as described previously^105^. Finally, Pearson correlations were performed between log-transformed power envelopes within each seed vertex and the rest of the brain to generate a PEC matrix for the seed-region. We then averaged the connectivity matrices to estimate rsEEG connectivity of the left IPL with the rest of the brain. Finally, the averaged connectivity maps for each frequency band were smoothed as commonly described in previous reports using a Gaussian kernel with a width of 5 x 5 x 5 mm full width at half maximum (Fig. 5B lower panels). We used the same local and DMN masks to extract average rsEEG measures. In addition, we computed global average across the cortex for each rsEEG measure.

### Cognitive measures

The Alzheimer’s Disease Assessment Scale-Cognitive Subscale (ADAS-cog) was administered to assess global cognitive function. The ADAS-cog, comprising 11 tasks assessing memory, language, praxis, and orientation, was conducted in a quiet environment by trained examiners over a 45 to 60-minute session. Tasks included word recall, object naming, command following, constructional praxis, ideational praxis, orientation, word recognition, and language comprehension, scored to reflect cognitive impairment severity. Total scores ranged from 0 to 70, with higher scores indicating greater impairment^108^. The Rey Auditory Verbal Learning Test (RAVLT)^109^ was administered to assess verbal memory and three components of interest were extracted: Immediate Recall, Delayed Recall, and Delayed Recognition. Participants were first given a list of 15 words to learn (List A) over 5 trials each. A second, distractor list of 15 words (List B) was then presented for the participants to learn. Immediate Recall of the initial list (List A) was then recorded as the number of words spontaneously recalled (range: 0-15, higher scores indicate better performance). The Delayed Recall score assesses the ability to recall the same list after a 20-minute delay without further exposure, evaluating long-term memory retention (range: 0-15, higher scores indicate better performance). The Delayed Recognition score involves presenting participants with a list of 30 words, including the original 15 words and 15 distractor words. Participants are required to identify which words were from the original 15 words, testing their recognition memory (range: 0-30, lower scores indicate worse recognition accuracy). The RAVLT total score is calculated by summing the scores from all components of the task (range: 0-150) with higher total scores reflecting better overall (global) verbal memory performance, indicating greater learning ability, retention, and recognition. Digit Span Task^110^ was administered to assess the working memory capacity of participants. The task included both Digit Span Forward and Digit Span Backward components. For the Digit Span Forward task, sequences of digits were read aloud at a rate of one digit per second, beginning with two digits and increasing incrementally by one digit per trial. Participants were required to repeat each sequence in the same order. This component primarily assesses immediate memory span and the capacity to hold and process information in a linear order. In the Digit Span Backward task, participants were asked to repeat the digit sequences in reverse order, following the same incremental sequence length. This task evaluates more complex working memory functions, as it requires not only retention of the information but also mental manipulation to reorder the digits. The tasks continued until the participant failed two consecutive trials of a particular sequence length. Performance was scored by the total number of trails completed, with higher scores indicating better performance (range: 0-16). The Animal Fluency Test was administered to evaluate the semantic memory and verbal fluency of participants. The test involved instructing participants to name as many animals as possible within 60 seconds. Participants were asked to name as many animals as they could think of in one minute, and their responses were recorded verbatim. The total number of unique animal names produced was counted, with repetitions, non-animal words, and proper nouns excluded from the final score. The task was scored by the total count of unique, valid animal names produced within the given time. This task is commonly used to assess executive control, lexical retrieval, and semantic memory^111^.

### Statistical Analyses

We performed cluster-based permutation independent sample *t*-test statistics to compare current density time series between AD and HC groups. First, we ran independent sample t-tests at each sample to determine significant time points. We then computed the length of adjacent significant time points and sum of t-scores for significant time points to determine (1) cluster size and (2) cluster magnitude in the main analyses, respectively. Following main analyses, we performed permutation t-tests (*n* = 1000) by randomly shuffling 50% of subjects across the groups (i.e., 50% of subjects shifted from AD to HC groups or vice versa) and determined cluster size and magnitude of significant adjacent time points at each iteration. Finally, we re-compute *p*-values of significant clusters in the main analyses by calculating the probability of their size and magnitude in the permutation analyses. A cluster in the main analyses is considered to survive permutation, and thus significant, only if both the size and magnitude of a given cluster is above 95% of all cluster sizes and magnitudes derived from permutation tests. For current density time series, we reported statistics both for main analyses as the average t-test scores and p values of the cluster and permutation analyses with the probability of permutation (Permutation p). For comparing average EEG activity (Fig. 2C violin plots in right panels) we performed independent sample t-test statistics.

For surface based analyses (Fig. 2D), we averaged current densities at time windows identified with the time series analyses described above, project them to MNI template and performed cluster-based Monte-Carlo permutations (n=1000) using the Field Trip “Fieldtrip function ft_sourcestatistics” implemented in brainstorm software with two-tailed independent t-tests and Cluster Alpha at *p* <=0.05. These surface statistics were performed as secondary analyses to get better insights on the spatial distribution of group differences.

To examine the relationships between EEG measures and cognitive function in the AD group, we performed Pearson product-moment correlations. We also assess bivariate relationships between potential confounders (age, education level, and IPL cortical-thickness) and our variables of interest (cortical excitability, network connectivity, and cognitive measures including ADAS-Cog, RAVLT-Total, Digit-Span Backward, and Fluency). To interpret the strength of the correlations, we classified the Pearson correlation coefficients as follows: correlations with absolute *r* < 0.1 were considered to indicate zero or no correlation, r > 0.1 and < 0.3 were considered weak, r > 0.3 and < 0.7 were considered moderate, and r > 0.7 were considered strong correlations. Following these preliminary analyses, we conducted hierarchical regression analyses to account for the potential confounding effects of age, education level, and cortical thickness on the relationships between EEG measures and cognitive outcomes. The hierarchical regression approach allows for the sequential entry of variables into the regression model, enabling the assessment of the unique contribution of EEG measures to cognitive outcomes after controlling for confounders. In the first step, we entered age, education level, and cortical thickness as control variables into the regression model. In the second step, we entered the EEG measure of interest (e.g., cortical excitability or DMN connectivity) into the model to determine its unique contribution to the cognitive outcome after accounting for the confounders. The change in R² (ΔR²) between Model 1 and Model 2 indicates the additional variance in the cognitive outcome that is explained by the EEG measure beyond that explained by age, education, and cortical thickness. We reported the standardized beta coefficients (β) for regression statistics. All statistical analyses, except surface based statistics, were performed using SPSS software version 21.0 (IBM Corp., Armonk, New York, USA).

