## Supplementary Material for "Neurophysiological signatures of default mode network dysfunction and cognitive decline in Alzheimer’s disease"

**Supplementary Table 1.** Participant demographics. Statistical comparisons of gender and handedness were performed using chi-square test, whereas all other comparisons were performed independent sample t-tests. MMSE: Mini Mental State Examination Test, MoCA: Montreal Cognitive Assessment, CDR-SB: Clinical Dementia Rating Sum of Boxes, ADAS COG: Alzheimer Disease Assessment Scale-Cognitive Subscale, RAVLT-Total: Rey Auditory Verbal Learning Test, Digit-Span Backward: Number of digit sequences recalled correctly in the reverse order. Fluency: The total number of unique animal names remembered correctly within 60 second.

|  | **AD**  **n=42** | **HC**  **n=40** | ***p*** |
| --- | --- | --- | --- |
| Age, years (std) | 70.95 (7.63) | 70.63 (6.41) | 0.84 |
| Female sex, no (%) | 18 (42.85) | 19 (47.5) | 0.67 |
| White race, no (%) | 37 (88.09) | 35 (87.5) | 0.93 |
| Education, years (std) | 17.38 (1.97) | 16.87 (3.00) | 0.57 |
| MMSE (std) | 25.87 (2.59) | 28.78 (1.19) | 0.00 |
| MoCA (std) | 20.61 (4.33) | 26.75 (2.22) | 0.00 |
| CDR-SB (std) | 2.5 (1.46) | 0.03 (0.13) | 0.00 |
| CDR-Global (std) | 0.59 (0.19) | 0.01 (0.08) | 0.00 |
| ADAS COG (std) | 14.94 (7.75) | 6.60 (3.56) | 0.00 |
| RAVLT-Total (std) | 51.90 (18.40) | 99.15 (18.22) | 0.00 |
| Digit-Span Backward (std) | 6.85 (2.59) | 8.72 (2.36) | 0.01 |
| Animal Fluency (std) | 13.61 (6.04) | 24.50 (4.59) | 0.00 |

**Supplementary Fig. 1. IPL excitability and network connectivity following M1 and Sham stimulations. A:** Averaged evoked responses from left IPL following M1 (upper left) and Sham-TMS (lower left) both for AD (red) and HC (Blue) groups. If present, auditory evoked potential (AEP) components were kept in Sham-data sets. Solid colored lines show group averaged current density time series (in z-scores) extracted from individualized e-field based local masks for IPL (orange shaded areas of the representative cortical surface in left panels). Shaded regions show inter-individual response variation with standard error of measurements (SE). Violin plots on the right panels show total amount of evoked current densities between 15-65ms following M1 (upper right panel) and Sham-TMS (lower right) for both groups. **B:** Cortical Maps for grand averaged current densities (in z-scores) between 15 and 65ms (upper and middle left panels) on a template brain model following Sham-TMS for AD (upper panels) and HC (middle panels) groups. Lower panels show statistical results of thresholded cluster-based permutation t-tests (cluster p<0.05) with hot colors indicating AD > HC and cold colors indicating AD < HC. Cortical maps for M1-TMS is provided in Fig.2.

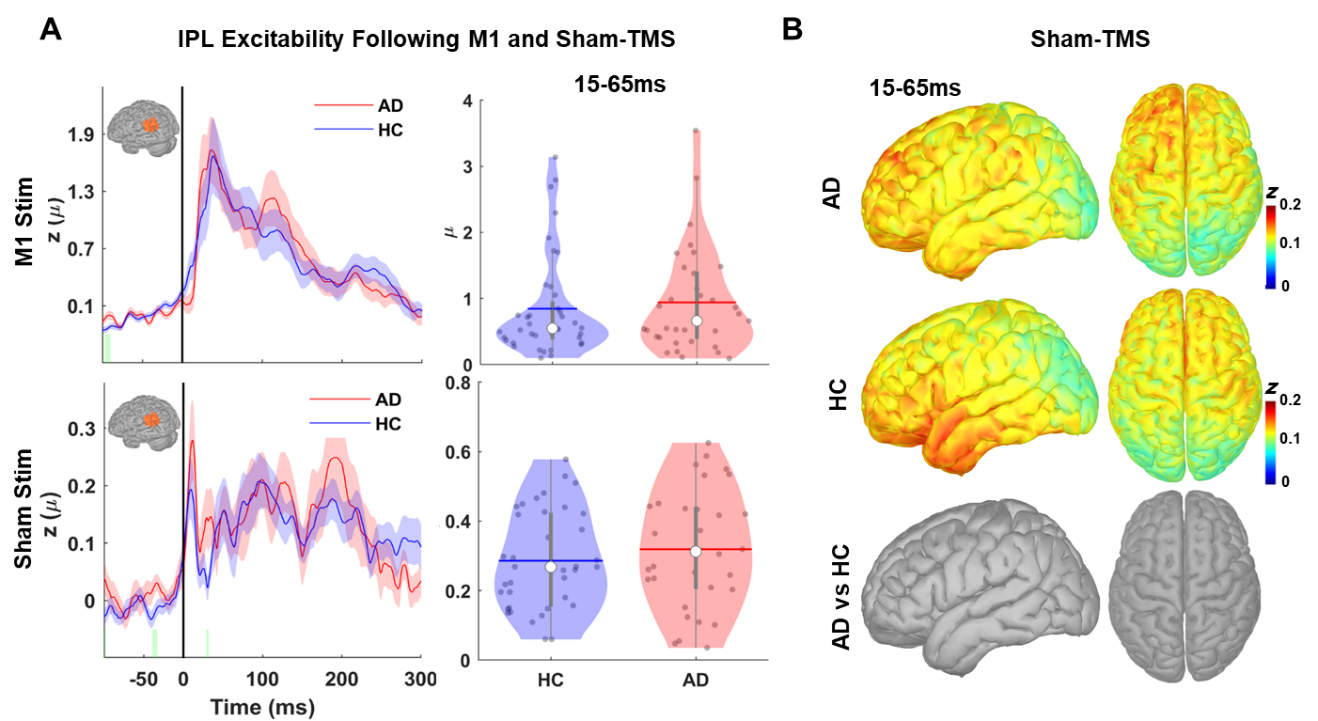

**Supplementary Fig. 2. Decreased Frontal DMN connectivity in AD. A:** Cortical maps of grand averaged current densities (in z-scores) between 125-175ms (right) on a template brain model following IPL-TMS for AD (upper panels) and HC (middle panels) groups. Lower panels show statistical results of thresholded cluster-based permutation t-tests (cluster p<0.05) with hot colors indicating AD > HC and cold colors in indicating AD < HC. **B:** Violin plots showing total amount of evoked current densities between 75-101ms following IPL-TMS for frontal DMN. **C**: Scatter plots with regression lines showing bivariate correlations of frontal DMN connectivity with Digit-Span (upper panel) and Fluency (lower panel). Color codes refer different temporal windows with red showing early (15-65ms) and blue showing late (125-175ms) activations. Correlation coefficients are provided for each regression line with asterisks indicating statistically significant correlations (p<0.05). Hierarchical regression analyses, controlling for age and cortical thickness, showed that late frontal DMN activations significantly predicted better Digit-Span (β=0.384, t=2.484, ΔR²=0.124, p=0.02) and Fluency (β=0.535, t=3.240, ΔR²=0.239, p=0.003) performance, explaining an additional 12% and 24% of unique variance.

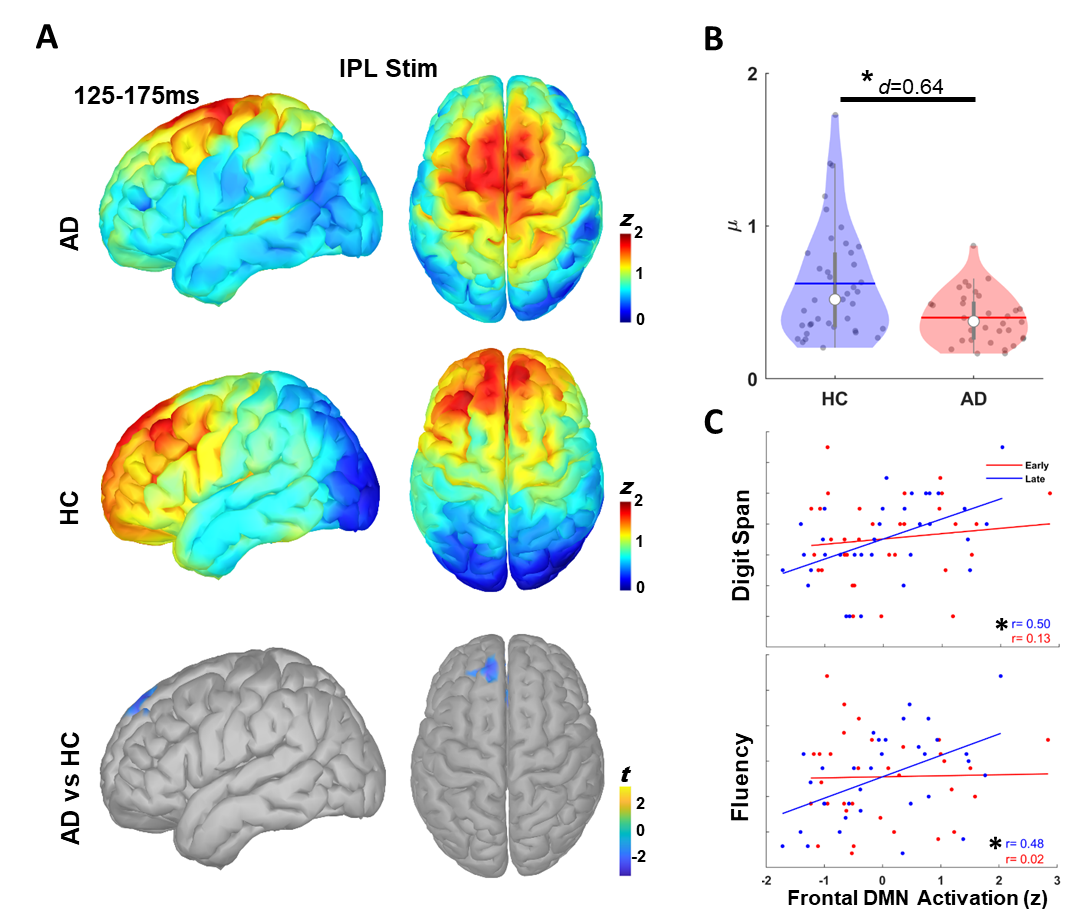

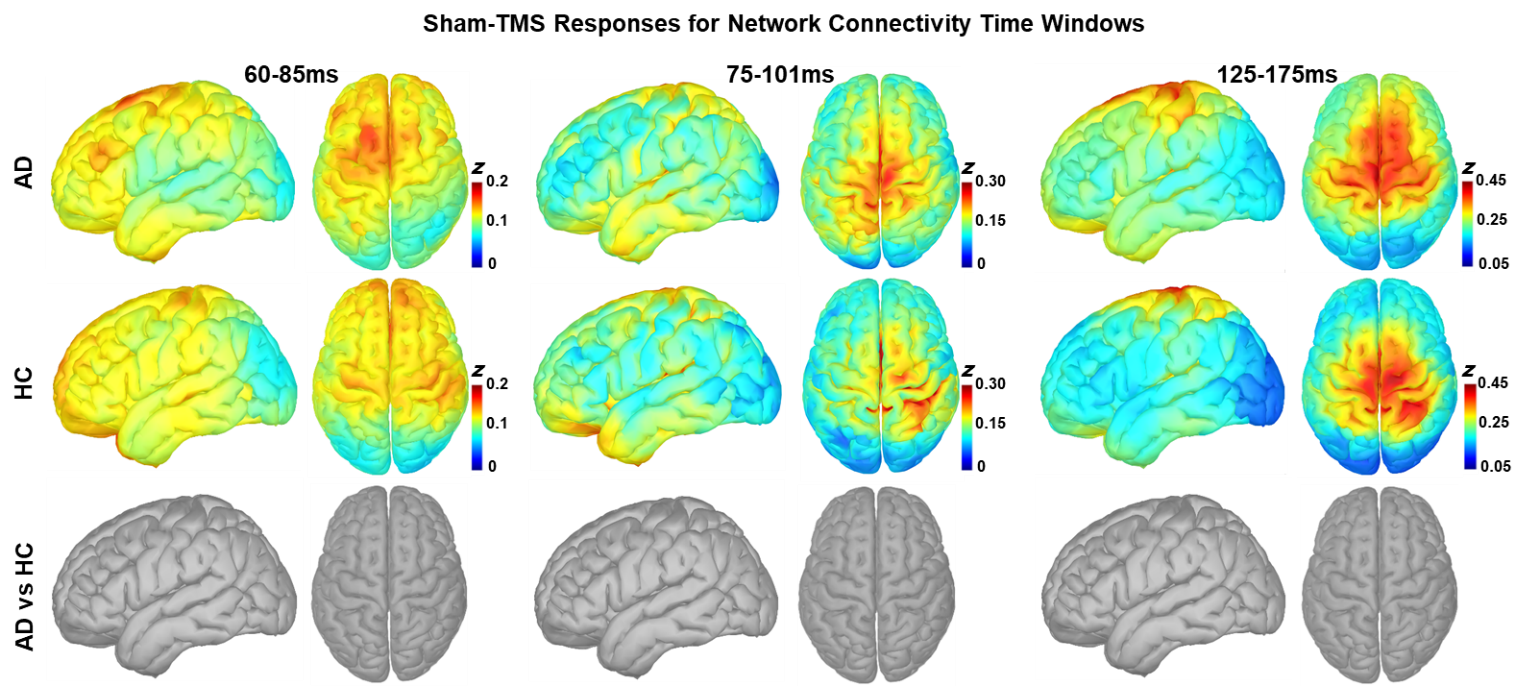

**Supplementary Fig. 3. Network connectivity with sham stimulation:** Cortical Maps for grand averaged current densities (in z-scores) between 60-85 (left panels), 75-101 (middle panels), 125-175ms (right panels) on a template brain model following Sham-TMS for AD (upper panels) and HC (middle panels) groups. Lower panels show statistical results of thresholded cluster-based permutation t-tests (cluster p<0.05) with hot colors indicating AD > HC and cold colors indicating AD < HC.

**Supplementary Table 2.** Bivariate Pearson correlations among control (Cortical Thickness, Age, Education), TMS-EEG and cognitive measures. Statistically significant r values are highlighted in red with * indicating p < 0.05 and ** indicating p < 0.01.

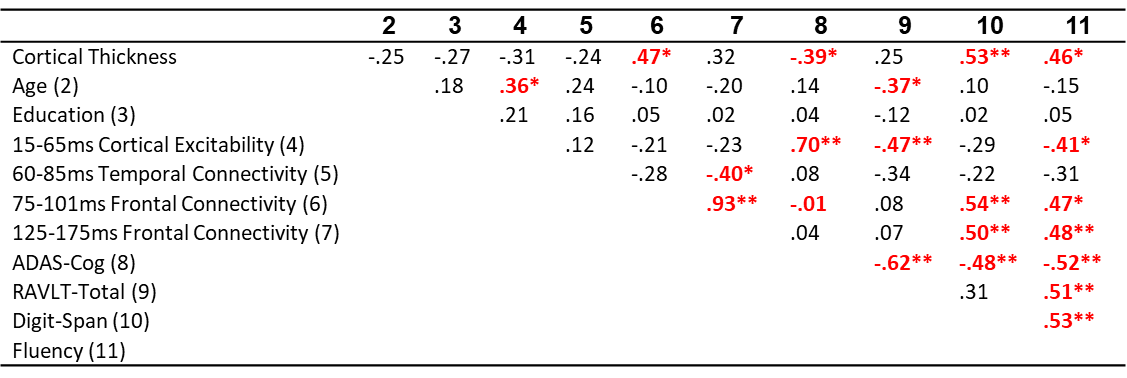

**Supplementary Fig. 4.** **Mediated Regression Diagrams. A:** Hierarchical regression models showing mediator effect of IPL cortical excitability on the relationship between cortical thickness and global cognition. Step-1 shows bivariate relationships between IPL cortical thickness, measured from local IPL maps, and ADAS-Cog scores in AD. Color coded circles (blue: cortical thickness and green: ADAS-Cog) show correlation variables with the degree of overlap in circles representing the amount of shared variance between them (Computed as R^2^). Step-2 illustrates the mediation model for cortical excitability. Black arrows between the variable blocks indicate bivariate relationships with color coded circles and correlation statistics (r for the strength and direction of the correlation, and p for the statistical significance). Color coded circles inside the mediation triangle shows results of the two-model hierarchical regression analyses. For the hierarchical regression, cortical excitability is entered as independent variable in the first model and cortical thickness is entered in the second model. The extent of over between red (cortical thickness) and green circles represents the amount of variance in ADAS-Cog explained by cortical excitability in the first model (R^2^ = 0.425), while the overlap between blue and green circles represents the amount of unique variance in ADAS-Cog explained by cortical thickness in the second model (R^2^ = 0.04) after controlling for the effects of cortical excitability. Regression statistics is shown with standardized β-coefficients (-0.211). Blue colored dashed arrow from cortical thickness to ADAS-Cog represents the reduced effects of cortical thickness on ADAS-cog as the significant bivariate relationship present in Step-1 became non-significant in Step-2 after controlling for cortical excitability. Step-3 illustrates mediation model for cortical thickness on the relationship between cortical excitability and ADAS-Cog. When cortical thickness entered as an independent variable in the first model, it explained 15% of variance in ADAS-Cog. Cortical excitability entered as an independent variable in the second model and explained an additional 34% of unique variance in ADAS-Cog. Red coded solid arrow from cortical excitability to ADAS-Cog with a significant β-coefficient indicates that cortical excitability is an independent predictor of ADAS-Cog after controlling cortical thickness. **B:** Hierarchical regression models showing mediator effect of Frontal DMN connectivity (for 75-101ms time window) on the relationship between cortical thickness and semantic memory with step-1 showing bivariate relationships between cortical thickness and Fluency, Step-2 illustrating the mediation model for frontal DMN connectivity and Step-3 illustrating mediation model for cortical thickness on the relationship between frontal DMN connectivity and semantic memory.

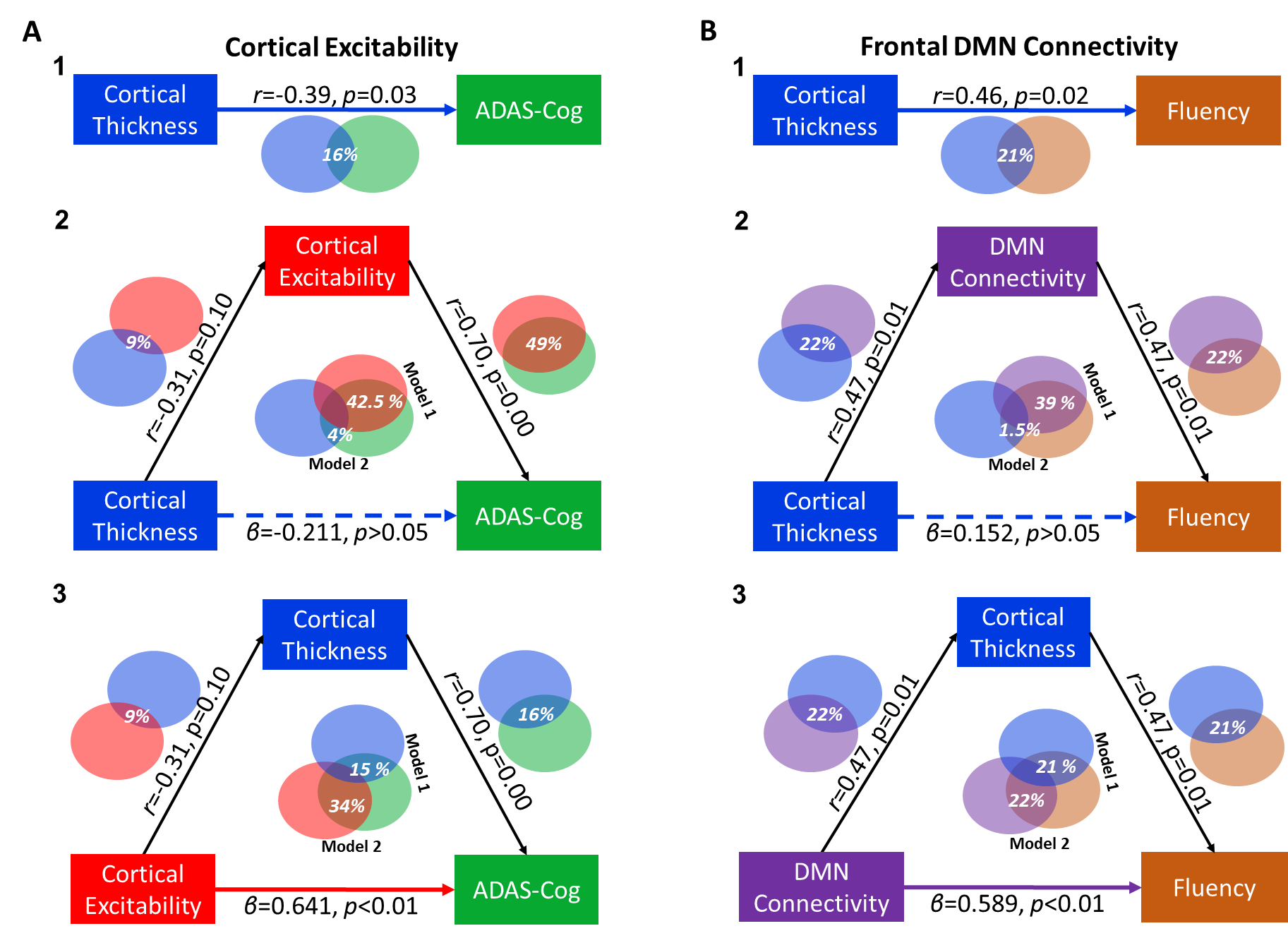

**Supplementary Fig. 5.** **Relationships between TMS-EEG measures of cortical excitability at IPL, network connectivity and cognitive functions in AD participants. A:** Scatter plots with regression lines in blue showing bivariate correlations of cortical excitability, extracted from M1 local cortical map following M1-TMS (left panels) and Sham-TMS (right panels) conditions, with cognitive functions. Red scatter plots and regression lines show correlations for IPL cortical excitability, already shown in Fig. 4 in the main text and provided again here for visual comparisons. **B:** Scatter plots with regression lines in blue showing bivariate correlations of network connectivity, extracted from frontal DMN node following M1-TMS (left panels) and Sham-TMS (right panels) for 75-101ms window, with cognitive functions. Red scatter plots and regression lines are already shown in Fig. 4 in the main text and provided again here for visual comparisons. Correlation coefficients are provided for each regression line with asterisks indicating statistically significant correlations (p<0.05).

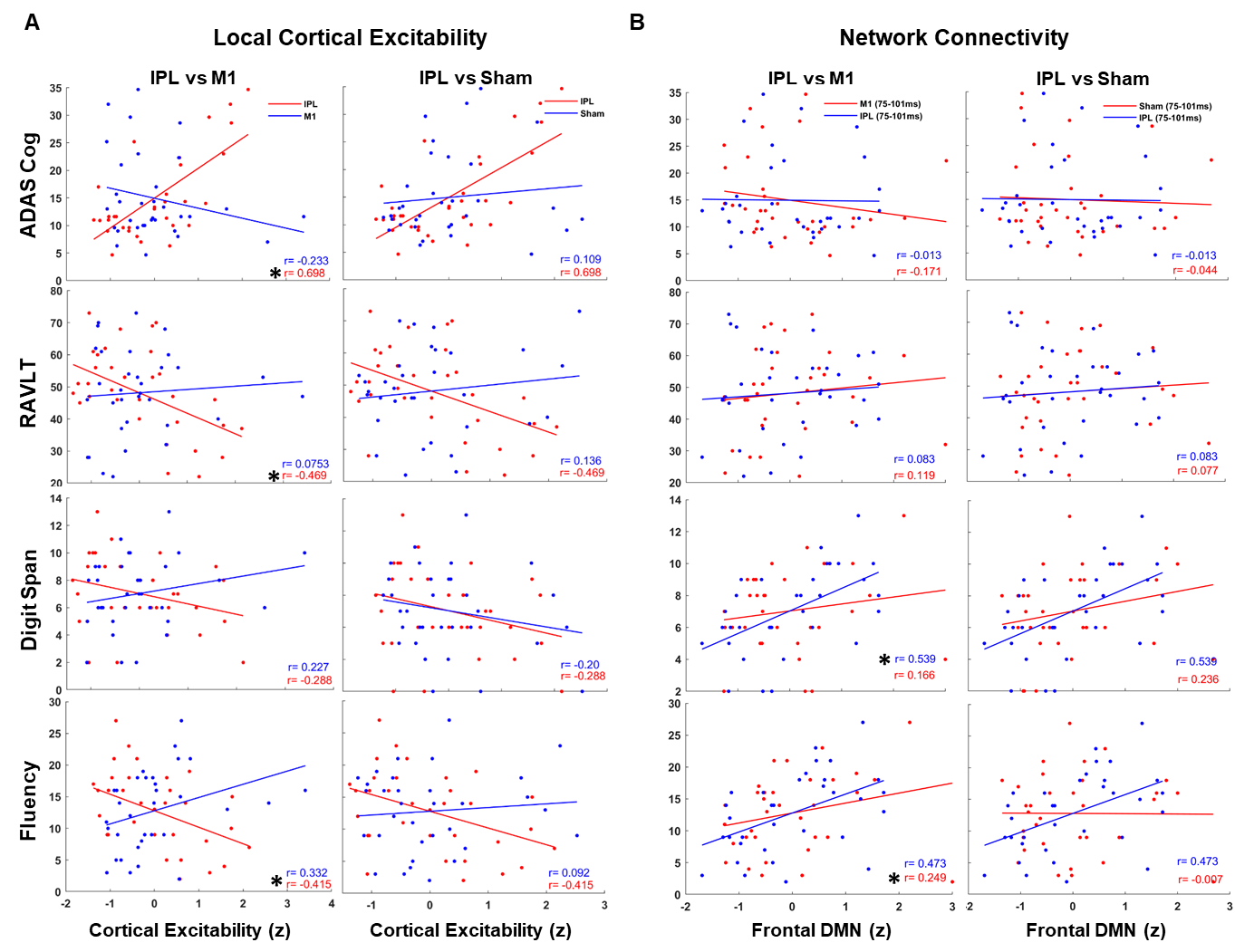

**Supplementary Fig. 6.** **Relationships between cortical excitability at Temporal-DMN, Precuneus-DMN and cognitive functions in AD participants. A:** Scatter plots with regression lines in red showing bivariate correlations of cortical excitability, extracted from temporal-DMN for early responses (15-65ms) following IPL-TMS compared with early responses following M1-TMS (blue plots in the left panels) and late responses (75-101ms) following IPL-TMS (blue plots in the left panels). **B:** Same illustrations in A for precuneus-DMN. Correlation coefficients are provided for each regression line with asterisks indicating statistically significant correlations (p<0.05).

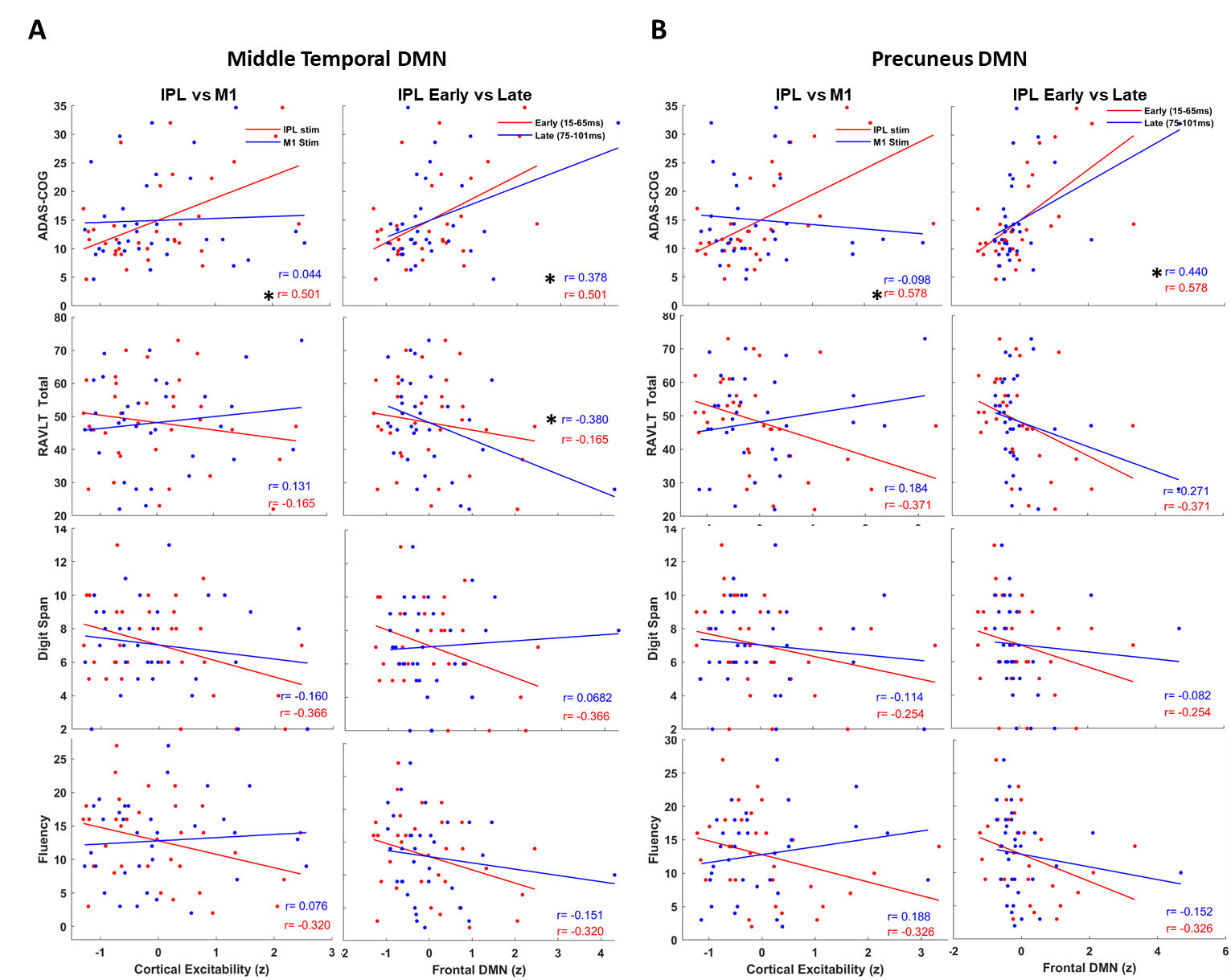

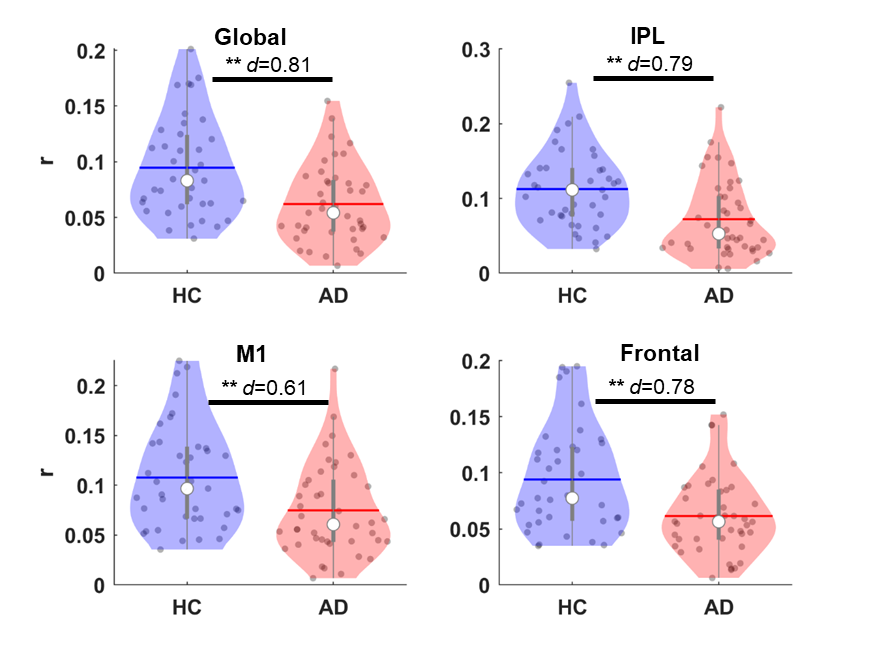

**Supplementary Fig. 7. Reduced beta-band neural synchrony in the AD.** Violin plots showing beta band neural synchrony averaged across the entire cortical space (global), within individualized masks of IPL, M1 and frontal-DMN both in healthy (blue) and AD (red) participants. * in upper violin plot denotes statistical significance at p<0.05 with corresponding effect size calculated using Cohens’ d.

**Supplementary Table 3.** Bivariate Pearson correlations among control (Cortical Thickness, Age, Education) and rsEEG measures of SPR and neural synchrony in the alpha band. Statistically significant r values are highlighted in red with * indicating p < 0.05 and ** indicating p < 0.01.

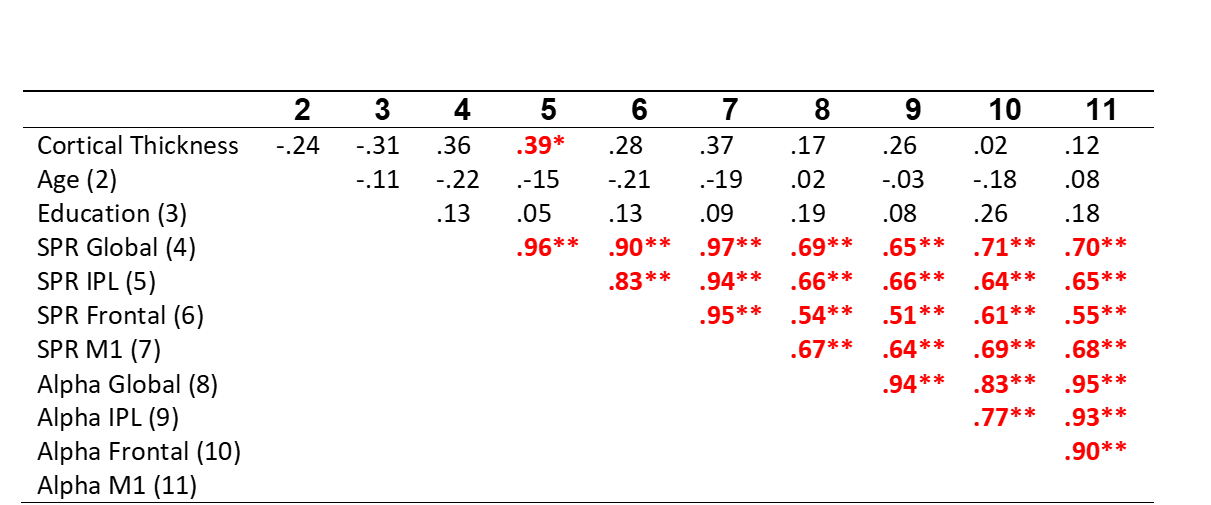

**Supplementary Fig. 8. Relationships between beta-band neural synchrony and cognitive functions in AD participants.** Scatter plots with regression lines showing bivariate correlations between neural synchrony in beta-band and cognitive functions in AD participants. Color codes refer different cortical regions with red showing average neural synchrony from local-IPL mask and blue showing from the frontal node of the DMN and black showing the average measure computed across the entire cortex (global). Correlation coefficients are provided for each regression line with asterisks indicating statistically significant correlations (p<0.05).

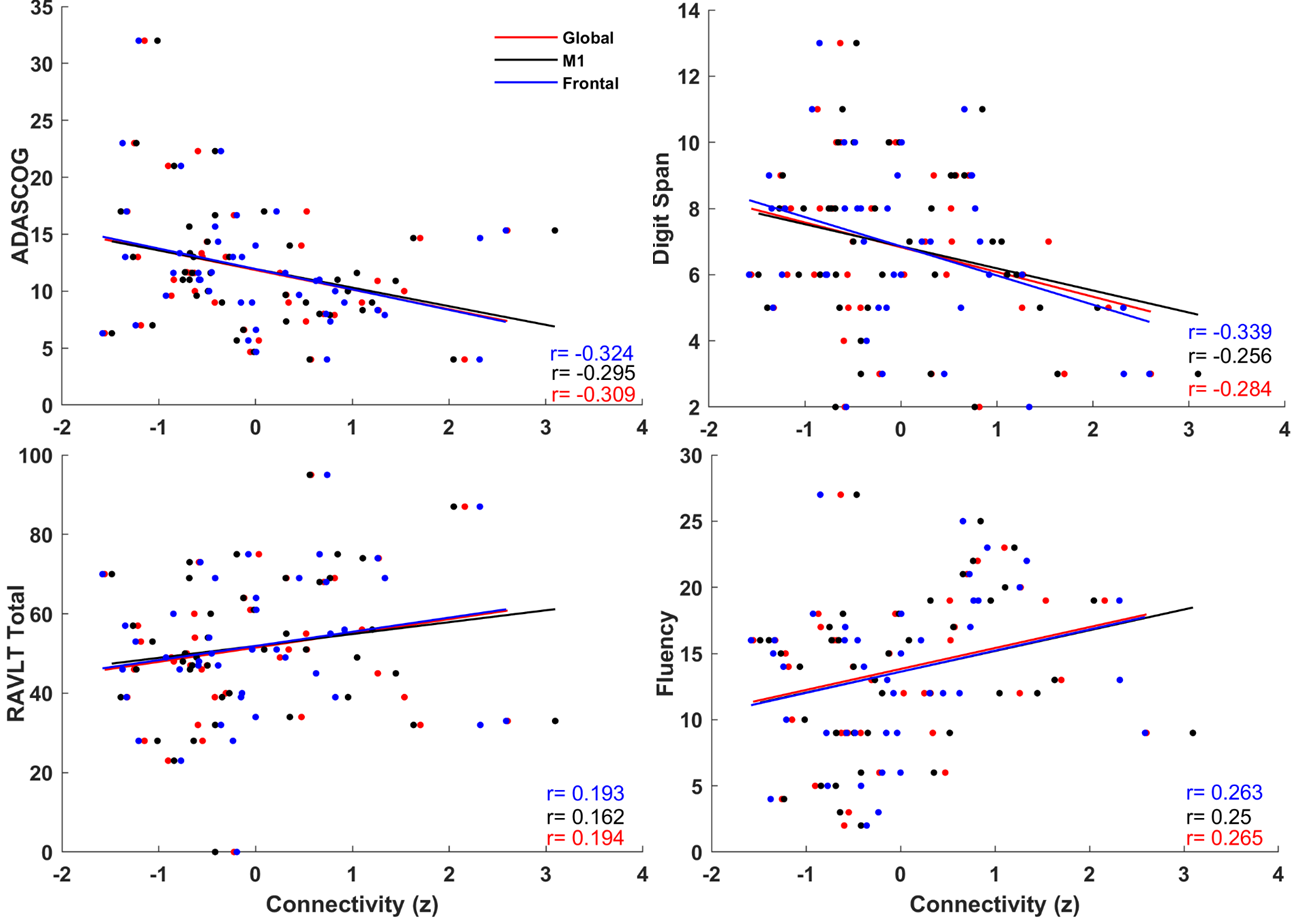

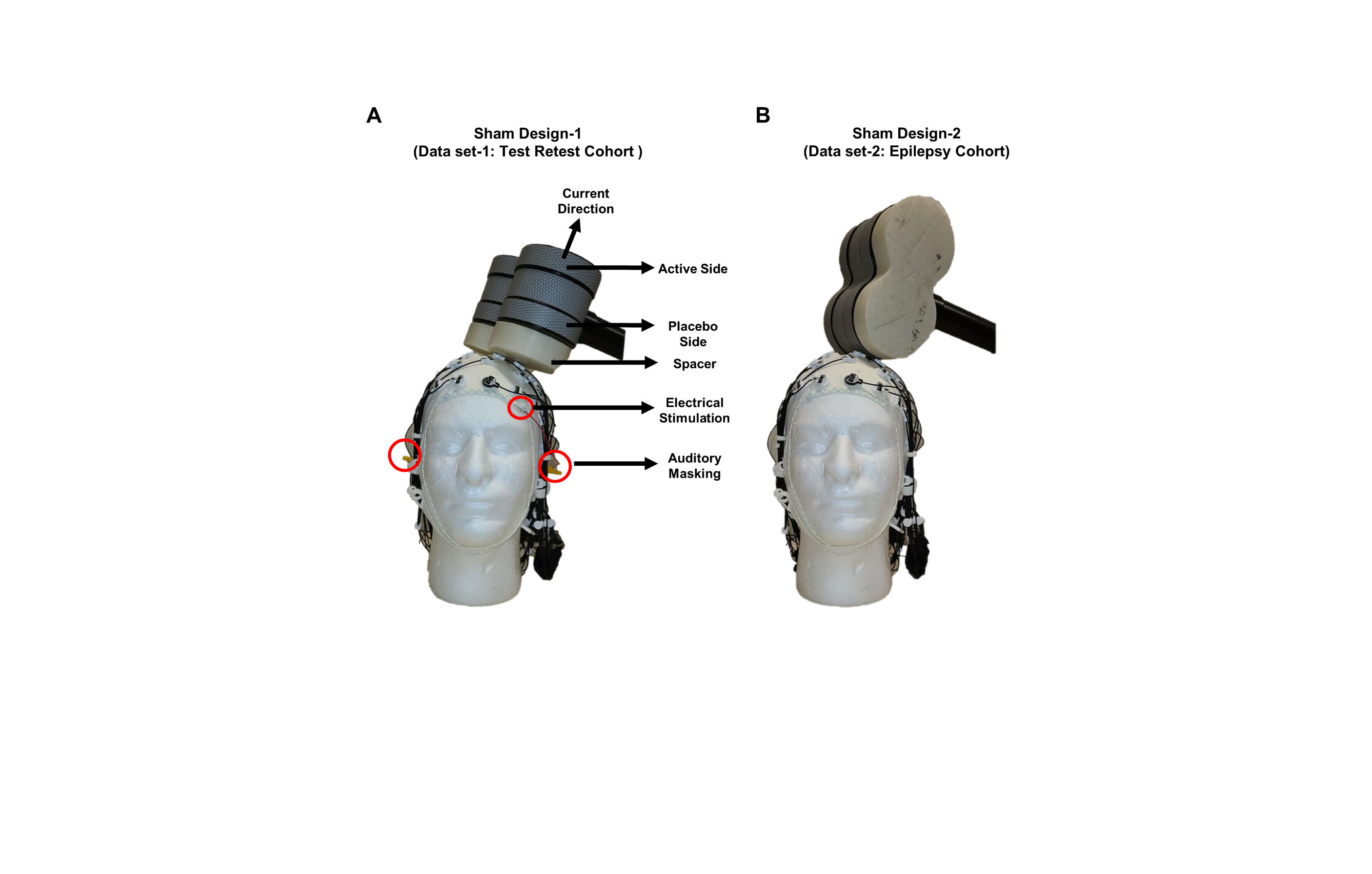

**Supplementary Fig. 9. Sham Protocol**. Sham TMS is delivered to L-M1 and TMS coil is flipped to the placebo side and 3cm spacer was placed under the coil. Auditory masking was used (red circles around the ears show earplugs) to minimize/eliminate AEPs and weak electrical stimulation is delivered to left frontal scalp region (red circle over the left eye shows location of surface electrodes) to mimic the current induced by the real TMS pulse.

**Supplementary Fig. 10. ICA decomposition from a representative subject**. **A:** Components identified as artifactual with visual inspection showing examples of eye blink, muscle activity, electrode noise and EKG components removed from EEG data. **B:** representative examples of neural components from occipital, sensorimotor, posterior/parietal and frontal regions.

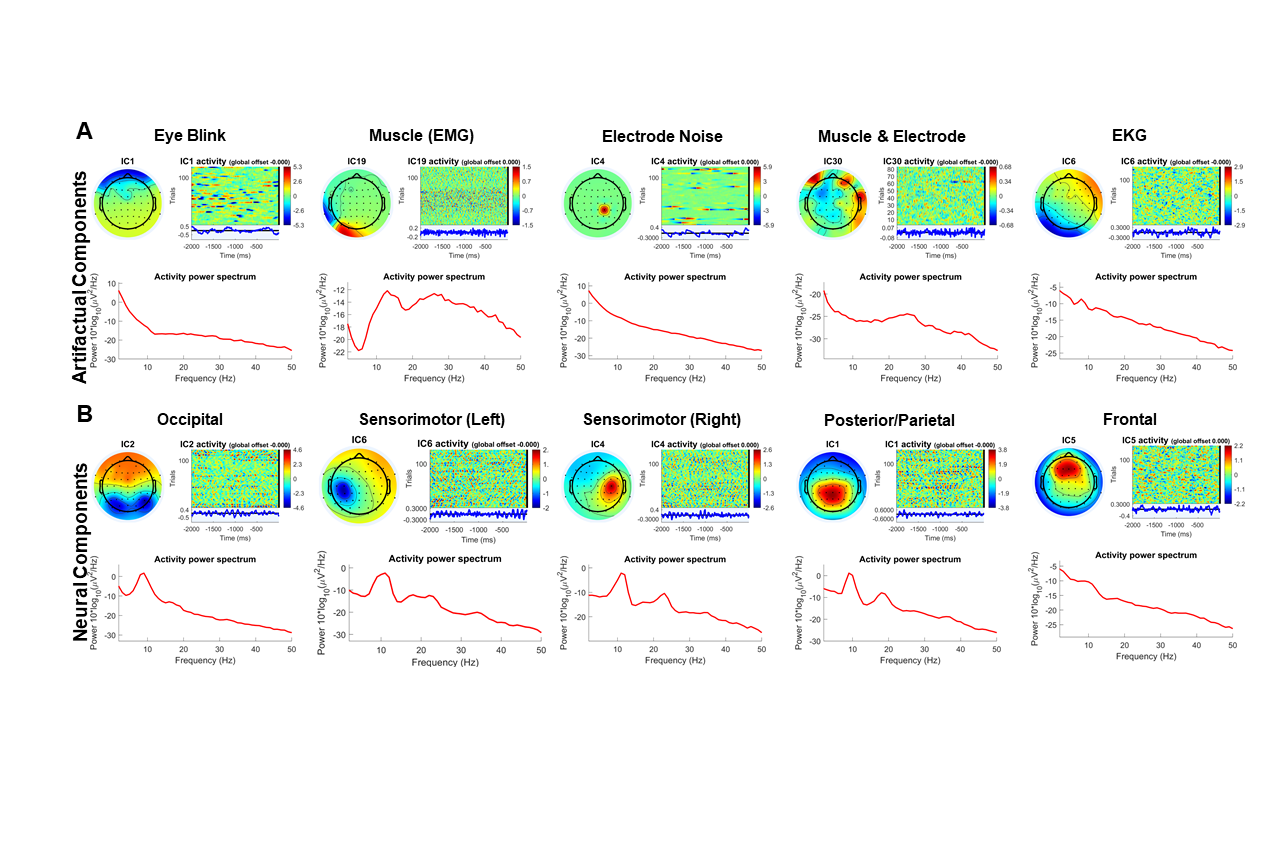

**Supplementary Fig. 11. Sham-informed ICA-based process for identifying and removing auditory evoked potentials. A:** Topographic distribution (left panels) and color-coded trial time series (right panels) of AEP component examples from TMS of IPL (upper panel) and sham-TMS of M1 (middle panel) for a representative participant. These two data sets were combined to perform a second round of ICA to identify and remove AEP from IPL data set. Note that both spatial topography and color-coded time series of individual trials were identical in both data sets and a uniform AEP component is perfectly aligned when both data sets were combined (lower panel). **B:** Average TEP topographies at different time points and TEP time series of IPL (red) and Sham (blue) data sets before (upper panel) and after (lower panel) removing the AEP component from combined data set. Note that TEP topographies of IPL after 70ms are almost identical to Sham topographies with AEPs present in both data sets (upper panel) but became distinct from Sham after removing AEPs. Also note that early TEP topographies of IPL at 30 and 70ms time points are identical regardless of the AEP condition (compare TEP topographies of IPL between upper and lower panels) suggesting that AEPs primarily contaminate TEPs at later components.

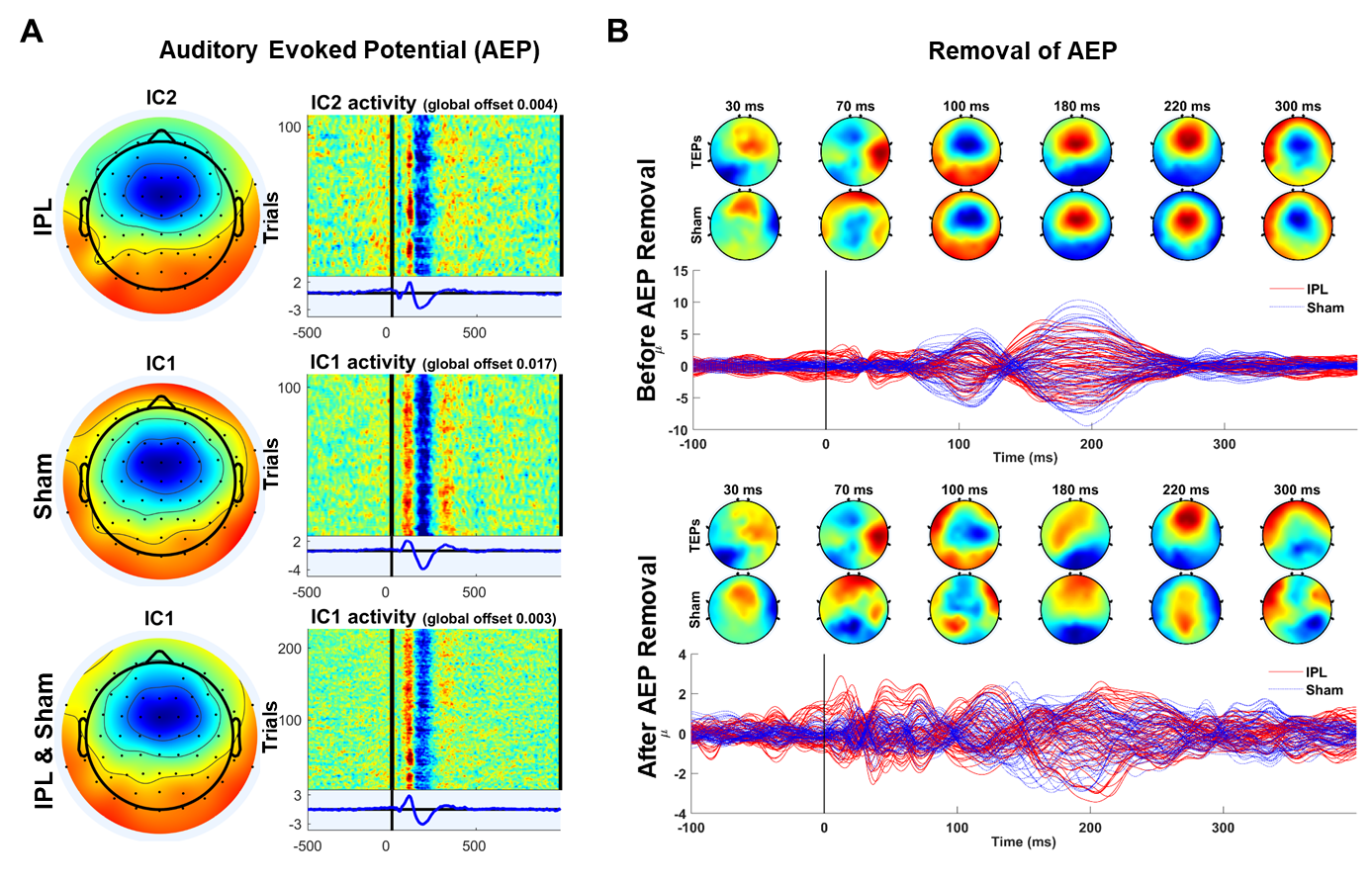

**IPL Trials**

**Sham Trials**

**Supplementary Fig. 12. Comparisons of resting motor threshold at M1, E-field, cortical thickness and scalp to cortex distance at IPL between AD and HC participants.** A: Individual E-field simulations for IPL (left panels) and M1 (right panels) are projected to template brain model to generate group average maps. Color bars show E-Field strength in units of V/m. **B:** Violin plots showing individual and group average data RMT (left upper), peak E-field strength (lower left), scalp to cortex distance (upper right), and average cortical thickness (lower right) at IPL. * in violin plots denotes statistical significance at p<0.05 using independent sample t-tests.

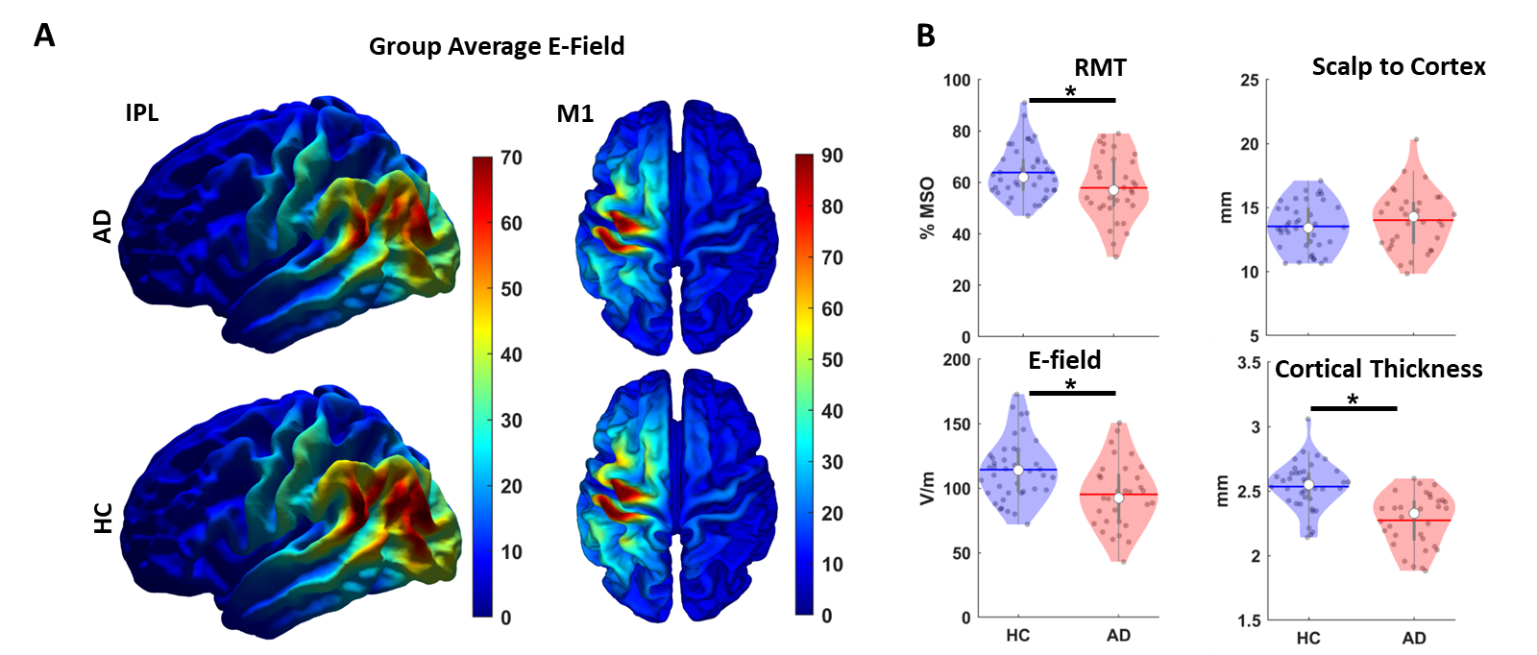
